# Host-adapted enzymatic deconstruction of acetylated xylan enables mutualistic colonization of monocot roots

**DOI:** 10.1101/2025.07.11.663926

**Authors:** Mathias Brands, Vicente Ramírez, Laura Armbruster, Ruben Eichfeld, Asmamaw Bidru Endeshaw, Pia Saake, Markus Pauly, Alga Zuccaro

## Abstract

Intracellular accommodation of mutualistic fungi in plant roots depends on selective remodeling of host cell walls while minimizing activation of plant immune responses. In this study, we identify a host-adapted enzymatic module in the root endophyte *Serendipita indica* that targets acetylated xylan, a major structural component of monocot cell walls. The glycoside hydrolase *Si*GH11 cleaves the xylan backbone and releases *O*-acetylated oligosaccharides, which are subsequently deacetylated by the XynE-like esterase *Si*AXE. These enzymes are co-expressed within a monocot-specific transcriptional program that is enriched in carbohydrate-active enzymes and sugar transporters. Their combined activity enhances enzymatic degradation and facilitates downstream hydrolysis by exo-xylanases, which reduces the production of apoplastic reactive oxygen species triggered by damage-associated molecular patterns. Functional analysis shows that overexpression of *Si*AXE promotes early root colonization, while deletion of the gene compromises fungal proliferation during later stages. These findings define a coordinated and immune-compatible strategy for host cell wall deconstruction that enables fungal adaptation and endophytic colonization of monocot roots.

**In Brief:** *Serendipita indica* utilizes a transcriptionally coordinated xylanase and esterase module to degrade acetylated xylan in monocot roots. This enzyme cooperation enhances substrate breakdown, suppresses immune responses, and enables endophytic colonization, illustrating how mutualistic fungi adapt saprotrophic enzymes for host-specific intracellular accommodation.

**Highlights:**

- Mutualistic fungal endophyte repurposes saprotrophic enzymes to enable monocot-specific intracellular root colonization
- Coordinated xylanase and esterase activity remodels host cell walls and dampens immune responses
- Expression of cell wall degrading enzymes is regulated by host species and colonization stage
- Results reveal fungal adaptation to monocot roots along the saprotrophy to symbiosis transition

## Introduction

Plants form complex associations with diverse microbial partners in the rhizosphere, ranging from mutualists to pathogens. Some of these microbes are highly specialized and can colonize the apoplast, a compartment encompassing the plant cell wall (CW) and extracellular space, which serves as a key interface for host–microbe interactions ^1,2^. The composition and architecture of the CW not only function as a physical barrier but also shape the biochemical landscape encountered by colonizing microbes. To adapt to this niche, both plants and microbes secrete proteins that remodel CW structure, including a broad suite of carbohydrate-active enzymes (CAZymes) ^3–5^. These include glycosyl transferases, polysaccharide lyases, enzymes with auxiliary activities, glycoside hydrolases (GHs), and carbohydrate esterases (CEs), which act on specific polysaccharides to mediate wall loosening, nutrient release, or niche establishment ^6,7^. In particular, microbial CAZymes must be adapted to the unique features of host species-specific CW polymers, such as the highly *O*-acetylated xylans found in monocotyledonous grasses ^8,9^.

The CW is a dynamic and complex matrix composed primarily of cellulose, hemicellulose, pectin, lignin, suberin, and CW-associated proteins ^10^. Hemicelluloses, such as xylan, xyloglucan, and glucomannan, form a network of β-1,4-linked backbones with diverse side-chain substitutions that influence polymer conformation, rigidity, and enzymatic accessibility. In monocot grasses, the predominant hemicellulose, xylan, is extensively modified. The β-1,4-xylosyl backbone is often acetylated at the *O-*2 and *O-*3 positions and can be further substituted with sugars such as arabinose, glucose, galactose, glucuronic acid and its methylated form ^11,12^. These modifications influence the conformation of the polymer and its interactions with other cell wall components, increasing wall rigidity and resistance to enzymatic degradation, while also establishing specific biochemical features that can be exploited by adapted microbial colonizers. Efficient degradation of such complex polymers requires coordinated enzymatic modules, typically composed of GHs, which cleave the polysaccharide backbone and its side chains, and CEs, which remove obstructive acetyl groups, either to facilitate further hydrolysis or to release acetate into the apoplast ^6,13^. The cooperation between these classes allows for stepwise deconstruction of decorated xylan, unlocking carbon sources and enabling fungal accommodation. The resulting oligosaccharides serve not only as nutrients but also as potential damage-associated molecular patterns (DAMPs) that can trigger plant immunity. Mutualistic fungi may counteract these responses by deploying additional effectors or enzymes to hydrolyze or sequester immunogenic fragments ^14–16^.

While microbial cell wall–degrading enzymes (CWDEs) are often associated with saprotrophy or pathogenesis, they are also present in mutualistic fungi, albeit in distinct repertoires. Ectomycorrhizal and arbuscular mycorrhizal fungi typically exhibit reduced CAZyme inventories, reflecting their predominantly extraradical growth and intracellular obligate biotrophic lifestyles, respectively. Mutualistic root endophytes with saprotrophic ancestry often retain or expand CWDE families, consistent with a more flexible trophic strategy ^17,18^. Although mutualistic fungi were traditionally thought to reduce enzymatic activity *in planta* to avoid triggering host defenses, recent evidence suggests that they may not loose these enzymes entirely. Instead, they might repurpose them through modular, tightly regulated expression programs. This suggests that enzyme-driven degradation can be compatible with the host immune system when it is carefully coordinated in space and time. Such strategies may help generalist endophytes adjust to hosts with different structures and immune responses by flexibly changing their metabolic activity. Understanding how mutualists integrate cell wall remodeling with immune evasion thus offers key insights into the evolution of symbiosis, the flexibility of fungal lifestyles, and the biochemical basis of host specificity. The mutualistic root endophyte *Serendipita indica* exemplifies this adaptive strategy, occupying an intermediate position along the saprotrophy-to-symbiosis continuum. Its CAZyme-rich genome includes expansions in xylan-degrading GH families, particularly GH10 and GH11, which encode canonical endo-β-1,4-xylanases specifically induced during colonization of monocot hosts ^19–21^. Comparative time-resolved transcriptomic analyses have revealed a striking host- and stage-specific upregulation of genes involved in CW degradation and nutrient acquisition, particularly during late-stage colonization of monocot grasses such as *Hordeum vulgare* (barley) and *Brachypodium distachyon* ^20–22^. The extensive repertoire of CWDEs in *S. indica* closely resembles that of saprotrophic fungi, suggesting that this symbiont dynamically adjusts its metabolic programs and intracellular colonization strategies in response to host cell wall composition and the physiological status of the host tissue, whether living or dead ^23^.

This plasticity aligns with its evolutionary background. *S. indica* and related Sebacinales are thought to have evolved mutualistic capabilities from saprotrophic ancestors, independently of pathogenic lineages, and typically lack the secondary metabolite gene clusters associated with virulence ^18,20,24^.

In pathogenic fungi, xylan-targeting GHs are well-established virulence factors and have been implicated in host infection across diverse species, including *Phytophthora*, *Verticillium*, *Magnaporthe*, *Sclerotinia*, and *Ustilago* ^25–29^. Similarly, microbial CEs that remove acetyl groups from xylan, commonly referred to as acetyl-xylan esterases (AXEs), have been associated with virulence and AXEs are found in CE families 1–7, 12, and 16 based on structural and biochemical classification ^13,30,31^. Despite extensive characterization in pathogenic fungi, the role of carbohydrate esterases targeting *O*-acetylated xylan in mutualistic plant fungal interactions and their potential cooperative activity with other CW modifying enzymes remains largely unexplored.

Here, using transcriptomic analysis across different host species, biochemical and immunogenic profiling of oligosaccharide products, and functional genetics in *S. indica*, we addressed the fundamental question of how mutualistic fungi adapt CW targeting enzyme systems to access host carbohydrates while supporting intracellular accommodation without compromising host integrity or triggering immune responses. We demonstrate a cooperative strategy involving a secreted xylanase *Si*GH11 and a XynE-like acetyl-xylan esterase *Si*AXE that together target *O*-acetylated xylan in monocot roots. Both enzymes are co-expressed in module 7, a monocot-specific transcriptional program enriched in CAZymes and transporters. *Si*GH11 releases acetylated xylooligosaccharides (XOS) which are then deacetylated by *Si*AXE, enhancing enzymatic efficiency and nutrient accessibility. *Si*AXE is required for efficient colonization, supporting a role in host compatibility and adaptation to monocot CWs. These findings uncover a tightly regulated enzymatic module that enables immune-compatible CW remodeling in mutualistic plant fungal interactions.

## Results

### *S. indica* activates a host- and stage-specific enzymatic program for xylan remodeling during monocot root colonization

To investigate the transcriptional response of *S. indica* during host colonization, we analyzed time-resolved RNA sequencing data collected at 1, 3, 6, and 10 days post inoculation (dpi) from interactions with the dicotyledonous model Arabidopsis thaliana and two monocot hosts, barley and *B. distachyon* ^21^. Weighted gene co-expression network analysis (WGCNA) of the monocot datasets revealed eight distinct gene modules composed of co-expressed *S. indica* genes showing coordinated temporal and host-specific expression (Extended Data Fig. 1A). These modules likely represent functional programs associated with adaptation to root environments, including host cell wall modification and nutrient acquisition. Among them, modules 4 and 7 were strongly enriched for genes encoding predicted secreted proteins and were preferentially induced during colonization of monocot hosts (Fig. 1A and Table S1). Notably, these two modules were either not detected or collapsed into broader, non-specific modules in the *A. thaliana* interaction, suggesting that the corresponding transcriptional programs are specific to monocot colonization. Module 7 showed the strongest induction for *in planta*–expressed genes at 10 dpi in *B. distachyon* and was significantly enriched for the Gene Ontology term hydrolase activity (GO:0016787). This module displayed an 11.5-fold enrichment in CAZymes relative to the genomic background, compared to only 0.5-to 2-fold enrichment in other modules (Fig. 1A–D, Extended Data Fig. 1A–B, Table S1). Most of the encoded CAZymes are predicted to act on host cell wall polysaccharides such as xylan, cellulose, and β-glucan (Fig. 1E). These features suggest that module 7 constitutes a specialized transcriptional program for extracellular remodeling of the host cell wall, specifically adapted to the structural complexity of monocot grasses roots during late-stage colonization.

**Figure 1.**
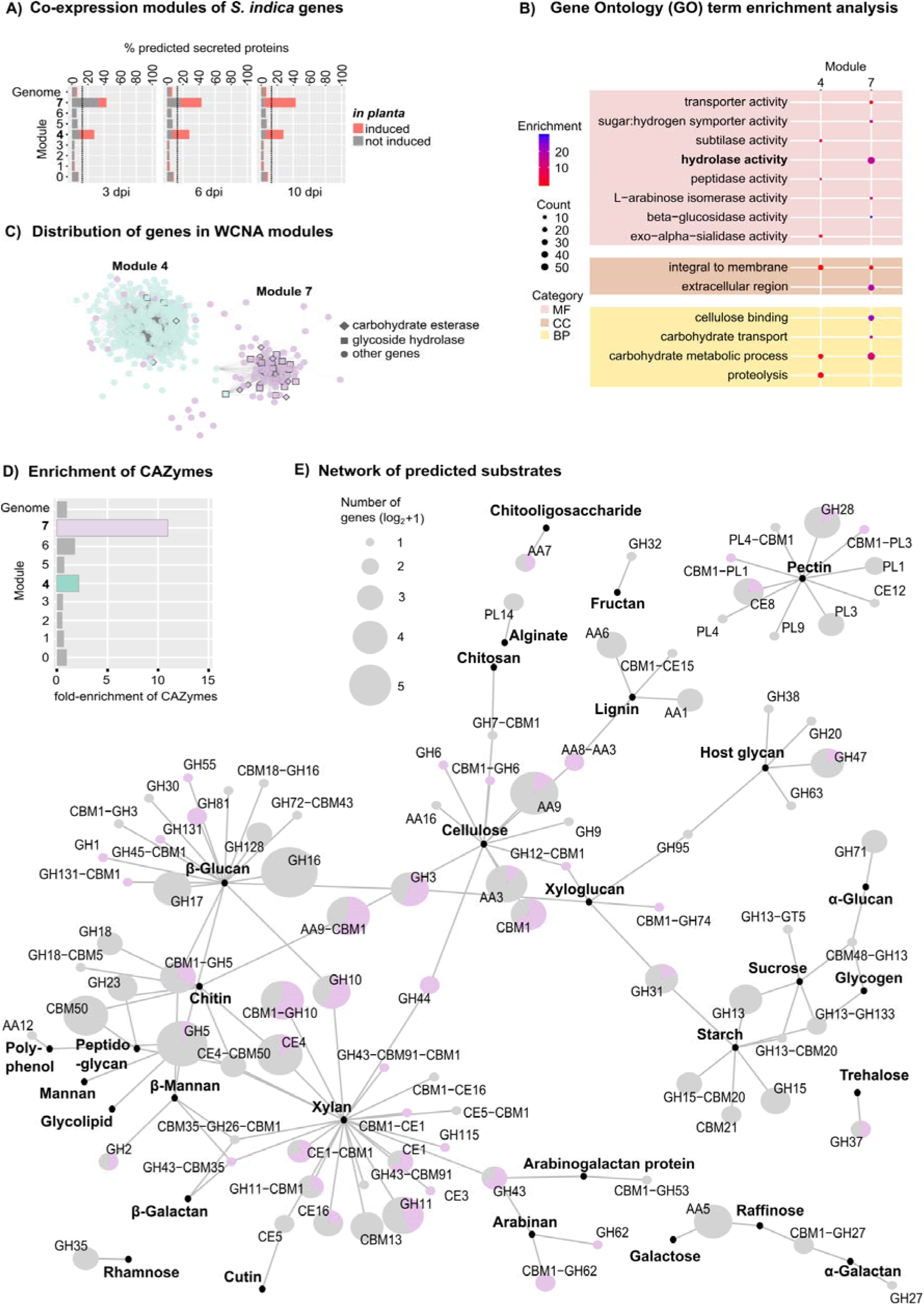
*S. indica* induces a monocot-specific co-expression module enriched in CAZymes during the late saprotrophic stage of root colonization. **A)** Weighted co-expression network analysis (WCNA) of RNA-Seq data from fungal transcripts collected at three time points post-inoculation (3, 6, and 10 dpi) reveals *in planta*–induced genes encoding predicted secreted proteins in modules 4 and 7. The dashed line indicates a two-fold enrichment compared to genomic background. **B)** Gene ontology (GO) enrichment analysis of molecular function (MF), cellular component (CC), and biological process (BP) categories for modules 4 and 7. Module 7 is notably enriched for hydrolase activity, including peptidase, glucosidase, and sialidase functions, and contains *SiGH11* and *SiAXE*. **C)** Composition of WCNA modules 4 and 7, highlighting the proportion of carbohydrate esterases (CEs) and glycoside hydrolases (GHs). **D)** CAZyme enrichment across WCNA modules 0–7 and relative to the whole genome. Enrichment is calculated as the ratio of CAZyme-encoding genes to total genes per module, normalized to the genome-wide CAZyme-to-gene ratio. **E)** Network of CAZymes annotated by predicted substrate class, based on domain composition. A total of 400 *S. indica* CAZymes were analyzed by counting individual genes (log_₂_+1 transformed) according to domain types (CBM, GH, CE, PL) or combinations thereof. Each gene was counted only once. Of the 181 genes in module 7, 78 are predicted CAZymes; the proportion of each CAZyme class in module 7 is shown in purple (pie charts). **Note:** CAZyme annotations are based on CAZyDB (http://www.cazy.org/), dbCAN3-HMMER, dbCAN3-sub ^4–5^, and manual curation by subfamily classification (see Table S1).

In addition to CAZymes, modules 4 and 7 also contain genes encoding proteins with predicted effector functions. These include the functionally characterized β-glucan-binding lectins *Si*FGB1 and *Si*WSC3, the ecto-5′-nucleotidase *Si*E5NT, the nuclease *Si*NucA, and several plant-responsive DELD proteins ^32–36^ (Table S1). The co-expression of these effectors with cell wall–modifying enzymes suggests that *S. indica* coordinates carbohydrate degradation with a secretome adapted for compatibility with the host apoplast.

Hydrolysis of acetylated xylan, the major hemicellulose in monocot grasses such as barley and *B. distachyon*, is a key determinant of host specialization in wood-decaying fungi ^37^. We hypothesized that the enrichment of CAZyme genes in module 7 reflects a targeted enzymatic strategy for xylan degradation during monocot colonization. To test this, we analyzed module 7 CAZyme expression at 10 dpi, the time point with peak expression in *B. distachyon*, alongside substrate predictions. Consistent with our hypothesis, a large proportion of module 7 CAZymes are predicted to target xylan, including 25 of 47 GHs and 12 of 32 CEs with xylan-degrading potential in the *S. indica* genome (Fig. 2A; Extended Data Fig. 1B). These expression patterns align with previous studies of microbial xylan-degrading enzymes and suggest both functional redundancy and transcriptional diversification in the fungal extracellular xylan degradation program ^4,13,38–41^.

**Figure 2.**
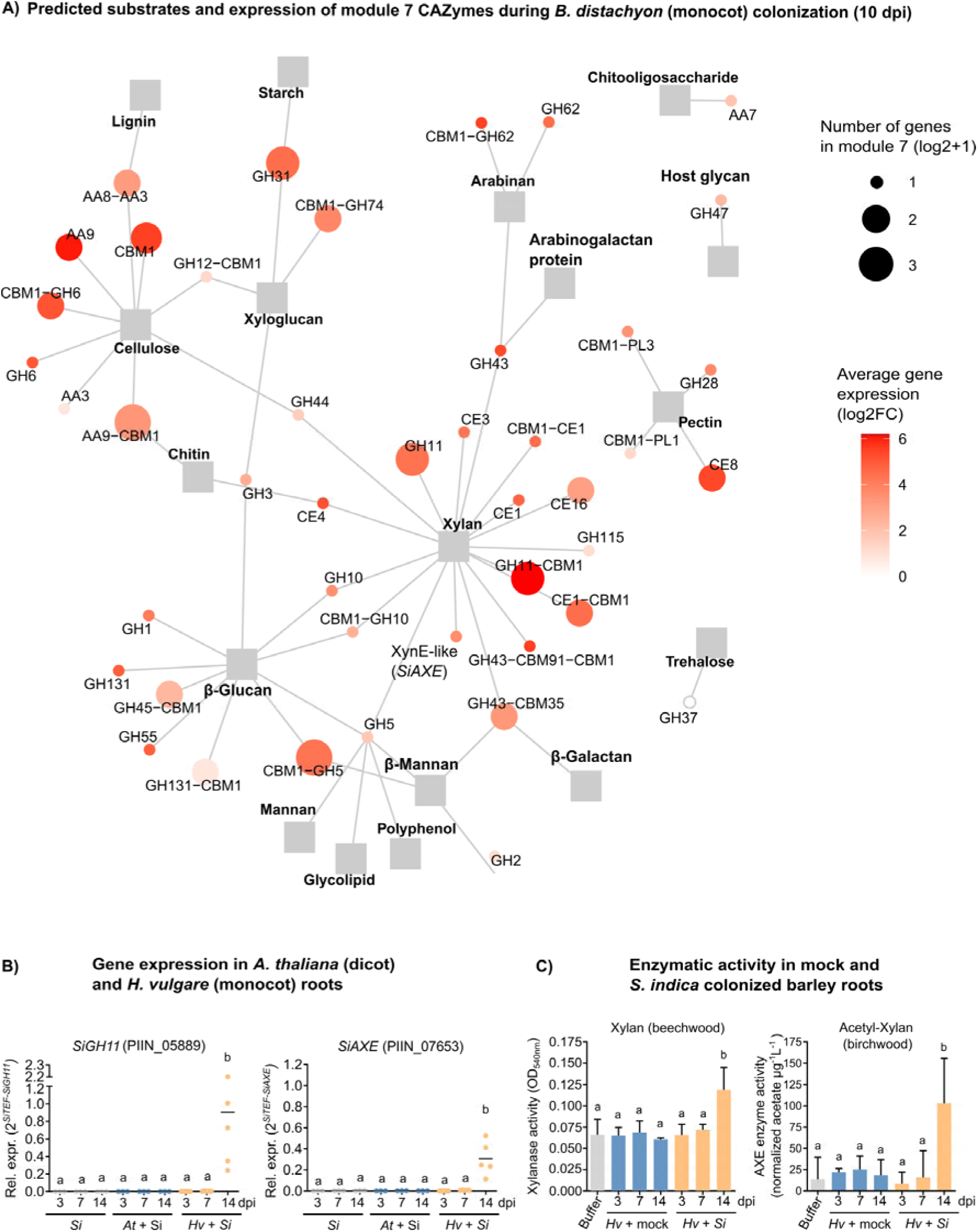
*S. indica* CAZymes of module 7 are induced during late-stage colonization of monocot hosts and contribute to acetyl-xylan degradation. **A)** Network of module 7 CAZymes annotated by predicted substrate class, based on domain composition. The module 7 CAZymes were analyzed by counting individual genes (log_₂_+1 transformed) according to domain types (CBM, GH, CE, PL) or combinations thereof. Each gene was counted only once. *S. indica* XynE-like (PIIN_07653; *SiAXE*) was added manually. Mean gene expression (log 2FC) was taken from *B. distachyon* colonized roots with *S. indica* at 10 dpi from previously published data ^21^. **Note:** CAZyme annotations are based on CAZyDB (http://www.cazy.org/), dbCAN3-HMMER, dbCAN3-sub ^4–5^, and manual curation by subfamily classification (see Table S1). **B)** Expression of *SiGH11* and *SiAXE* in *S. indica* mycelium from axenic culture or during colonization of the dicot host *Arabidopsis thaliana* or the monocot host *H. vulgare*. Both genes are strongly induced at 14 dpi in barley roots, corresponding to the late saprotrophic phase of colonization, consistent with previously published data ^22^. **C)** Enzymatic activity of root protein extracts from mock-inoculated and *S. indica*-inoculated barley at 3, 7, and 14 dpi. Xylanase activity was assayed using beechwood xylan as a substrate and quantified by the DNSA method (reducing sugars, OD_540_nm). Acetyl-xylan esterase (AXE) activity was measured using birchwood acetyl-xylan and quantified as released free acetate (buffer-subtracted). Protein extraction buffer served as a negative control. Different letters indicate significant differences (ANOVA followed by post hoc Tukey’s test, p ≤ 0.05).

To gain mechanistic insight into the function of the module 7 CAZyme program during monocot root colonization, we selected two co-induced secreted enzymes for functional characterization. *Si*GH11 (PIIN_05889) is a predicted GH family 11 endo-β-1,4-xylanase previously identified as a marker of the cell death–associated phase of colonization in barley^22^. *Si*AXE (PIIN_07653) is a single-copy, XynE-like acetyl-xylan esterase with structural similarity to an enzyme from the hemicellulolytic bacterium *Prevotella bryantii*, found in a characterized xylan-utilization cluster ^42^. Quantitative PCR confirmed that both genes are strongly induced at 14 dpi in barley roots, while expression remains low in *A. thaliana* and in axenic culture (Fig. 2B), consistent with prior transcriptomic data ^22^. Supporting their functional relevance *in planta*, xylanase and acetyl-xylan esterase activity was elevated in protein extracts from colonized barley roots at this time point (Fig. 2C).

Together, these findings identify xylan degradation as a central feature of a transcriptional program that supports cell wall remodeling and intracellular colonization in monocot roots. This enzymatic activity is embedded within a broader adaptive response that coordinates host species–specific secretion of cell wall–modifying enzymes and effectors. During this late colonization stage, *S. indica* primarily occupies dying or senescing root cells, reflecting a shift toward a saprotrophic-like growth mode.

### *Si*GH11 functions as an endo-1,4-β-xylanase

To characterize *Si*GH11 enzymatic activity, we expressed and purified the recombinant protein and tested its substrate specificity. *Si*GH11 contains a C-terminal carbohydrate-binding module (CBM1) and two conserved glutamate residues in its catalytic center, consistent with canonical GH11 xylanases (Extended Data Fig. 2A-C). The enzyme showed strong activity on azo-xylan (Fig. 3A), beechwood xylan, and linear XOS, which represent model substrates of unsubstituted β-1,4-xylan backbones (Fig. 3B). *Si*GH11 ultimately released xylobiose (Xyl2) and xylotriose (Xyl3). Co-incubation with the exo-xylanase *Sr*GH43 from *Streptomyces rochei*, which cleaves XOS from the non-reducing end, further digested these products to xylose (Fig. 3B), indicating that complete depolymerization requires cooperation with exo-acting enzymes. *Si*GH11 showed no detectable activity on unrelated microbial or plant cell wall polysaccharides such as, chitin, glucomannan or cellulose (Extended Data Fig. 2D), supporting substrate specificity for β-1,4-xylan.

**Figure 3.**
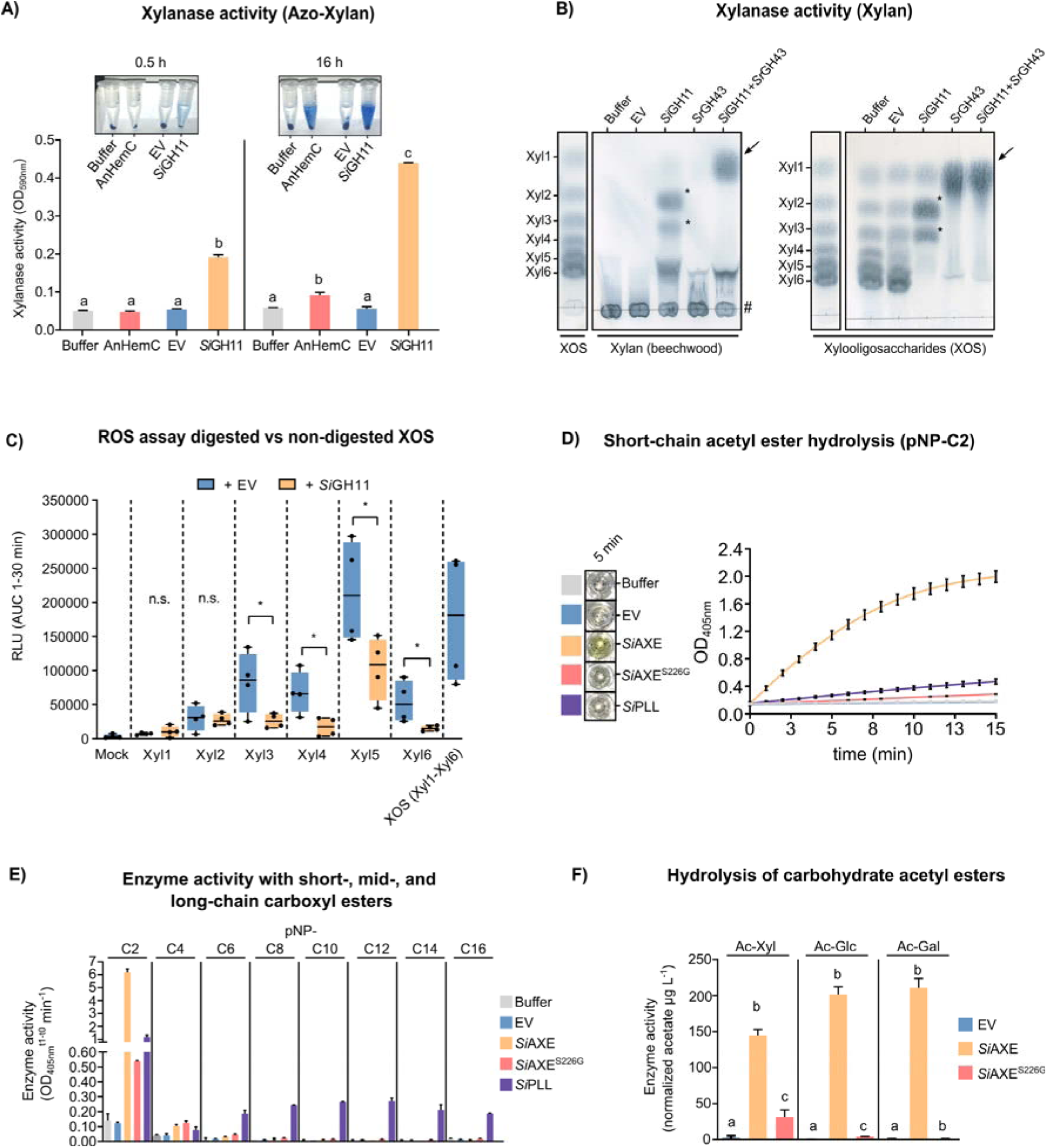
Biochemical activity of the *S. indica* endo-xylanase *Si*GH11 and acetyl-esterase *Si*AXE. **A)** Recombinant *Si*GH11 exhibits xylanase activity on azo-xylan (1% w/v). Substrate was incubated for 0.5 or 16 h with buffer, empty vector (EV) control, positive control (*Aspergillus niger* Hemicellulase C, AnHemC), or recombinant *Si*GH11. Activity was visualized as a shift in absorbance (OD_590_nm) following ethanol extraction of the supernatant. Different letters indicate significant differences (ANOVA + Tukey’s post hoc test, p ≤ 0.05). **B)** Thin-layer chromatography (TLC) analysis of xylanase activity by *Si*GH11 alone or in combination with exo-xylanase *Sr*GH43, using beechwood xylan (left) and linear unsubstituted xylooligosaccharides (XOS; right) as substrates. XOS mixtures served as size markers. Xylobiose (Xyl2) and xylotriose (Xyl3) bands are marked with asterisks; longer oligomers with a pound sign (#). Substrate specificity of *Si*GH11 was validated on a range of plant polysaccharides (see Fig. S3). **C)** Reactive oxygen species (ROS) burst in barley root segments triggered by XOS (Xyl1–Xyl6) before and after *Si*GH11 treatment (16 h incubation). *Si*GH11 digestion reduces ROS induction by longer XOS (Xyl3–Xyl6). A mixture of Xyl1–Xyl6 served as a positive control. Asterisks indicate significance (Student’s t-test; ns = not significant, *p* ≤ 0.05). AUC, area under the curve (see Figure S4 for kinetics). **D)** Acetyl-esterase activity of recombinant *Si*AXE with para-nitrophenyl acetate (pNP-C2), a short-chain artificial substrate. Ester hydrolysis is indicated by a colorimetric shift at OD_405_nm. *Si*AXE is highly active on pNP-C2. **E)** Substrate specificity of *Si*AXE and related enzymes tested with pNP esters of increasing chain length (C2–C16). *Si*AXE shows strong activity only toward pNP-C2. The S226G catalytic mutant has reduced activity. By contrast, *Si*PLL displays broad substrate specificity, consistent with lipase activity (see Fig. S6). **F)** *Si*AXE activity on acetylated carbohydrates: acetyl-xylan (Ac-Xyl; beechwood), glucose-pentaacetate (Ac-Gluc), and galactose-pentaacetate (Ac-Gal). Activity was quantified after 1 h by measuring free acetate release (buffer-subtracted). *Si*AXE is active on all three substrates, while the *Si*AXE^S226G^ mutant shows diminished but detectable residual activity. Different letters indicate statistical significance (ANOVA + Tukey’s post hoc test, p ≤ 0.05).

To assess whether *Si*GH11 modulates the immunogenicity of xylan-derived oligosaccharides *in planta*, we measured apoplastic ROS production in barley roots as a proxy for DAMP-triggered immune responses ^43^. Treatment with commercial XOS (Xyl1 to Xyl6) elicited varying levels of ROS, with Xyl5 triggering the strongest response. Pre-digestion of XOS with *Si*GH11 significantly reduced the ROS burst elicited by Xyl3 to Xyl6 (Fig. 3C; Extended Data Fig. 2E), suggesting that *Si*GH11 degrades immunogenic oligosaccharides into shorter, less active fragments.

These results demonstrate that *Si*GH11 is a functional endo-1,4-β-xylanase that degrades both polymeric xylan and linear XOS, ultimately producing Xyl2 and Xyl3, and that its activity attenuates the immune-stimulatory potential of xylan-derived oligosaccharides in the apoplast.

### *Si*AXE functions as a xylan acetyl esterase

To investigate the enzymatic mechanism underlying acetyl group removal from xylan, we functionally characterized *Si*AXE. This enzyme belongs to the SGNH hydrolase superfamily and contains a conserved Ser–His–Asp catalytic triad, along with an N-terminal CBM-like domain. Predicted structural modeling revealed homology with characterized microbial acetyl-xylan esterases (Extended Data Fig. 3A–C). The *Si*AXE enzyme efficiently hydrolyzed para-nitrophenyl acetate (pNP-C2), a standard proxy for short-chain acetyl esterases, but showed no activity on longer-chain pNP esters (Fig. 3D–E). In contrast, the promiscuous *S. indica* lipase *Si*PLL (PIIN_06162), used as a positive control, exhibited broad activity across short-, mid-, and long-chain esters and lipids (Fig. 3E; Extended Data Fig. 4). *Si*AXE also deacetylated carbohydrate-based substrates including acetyl-xylan, glucose-pentaacetate, and galactose-pentaacetate (Fig. 3F). Substitution of the predicted catalytic serine (S226G) resulted in a marked reduction in esterase activity across all tested substrates (Fig. 3D–F), confirming its essential role in catalysis.

These findings demonstrate that *Si*AXE is a carbohydrate esterase with specificity for *O*-acetylated sugars, consistent with its predicted role in deacetylating xylan-derived oligosaccharides *in planta*.

### *Si*GH11 and *Si*AXE cooperatively deconstruct native acetylated xylan from barley roots

Given their co-induction during late-stages colonization of barley and *B. distachyon*, we hypothesized that *Si*GH11 and *Si*AXE act cooperatively to deconstruct acetylated xylan in monocot root cell walls. To test this, we treated delignified alcohol-insoluble residue (dAIR) from barley roots with recombinant *Si*GH11, *Si*AXE, the exo-xylanase *Sr*GH43, or their combinations (Fig. 4). *Si*AXE^S226G^ served as the catalytically inactive control.

**Figure 4.**
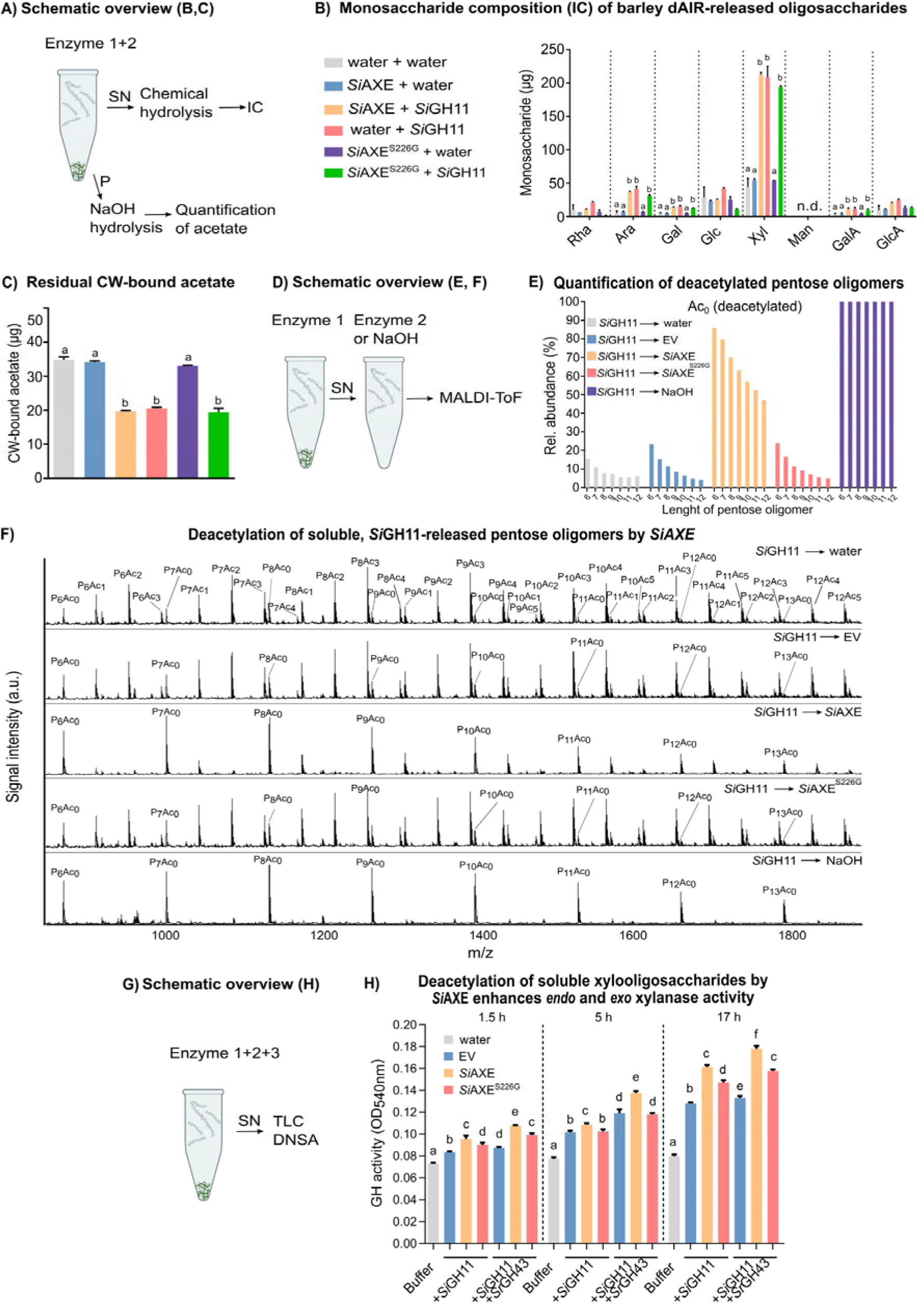
*Si*GH11 and *Si*AXE cooperatively deconstruct acetylated xylan from barley root hemicellulose. **A)** Schematic overview of the experiment in B and C. Barley delignified alcohol insoluble residue (dAIR) was treated with *Si*AXE or *Si*GH11 or a combination and the supernatant (SN) and pellet (P) used for analysis of monosaccharides and residual CW-bound acetate, respectively. **B)** The supernatant of dAIR enzyme digests was used for quantification of monosaccharides via ion chromatography after chemical hydrolysis. **C)** The pellets of the dAIR enzyme digests was treated with NaOH to hydrolyze all residual CW-bound acetate which was subsequently quantified as free acetate. **D)** Schematic overview of the experiment in F and G. Barley dAIR was first treated with *Si*GH11 and the supernatant, containing *Si*GH11-released oligosaccharides, harvested treated with *Si*AXE or controls and then analyzed with MALDI-ToF. **E)** Relative abundance of fully deacetylated oligomers (P6–12/Ac0) following enzymatic treatment or controls. Incubation with *Si*AXE or NaOH increases the proportion of non-acetylated pentose oligomers. NaOH treatment leads to near-complete deacetylation. Quantification of partially acetylated oligomers (Ac1–Ac6) confirmed *Si*AXE- and NaOH-dependent deacetylation (see Fig. S7A-F). **F)** MALDI-ToF spectra of oligosaccharides after enzymatic treatments shown in D and E. Peaks are annotated by the number of pentose units (P6–12) and acetylations (Ac0–6). **G)** Schematic overview of the experiment depicted in H. Barley dAIR was incubated with *Si*GH11, *Si*AXE and *Sr*GH43, alone or in combinations and the supernatant harvested for TLC analysis of soluble oligosaccharide reaction products and DNSA assay for quantification of reducing end sugars. **H)** Quantification of reducing ends in hydrolysis reactions using the DNSA assay. Samples were taken at 1.5h, 5h and 17 h to monitor GH activity. For TLC analysis, see Fig S7G-H. Reducing sugar release was measured as OD_540_nm shift. Different letters indicate statistically significant differences (ANOVA + Tukey’s post hoc test, p ≤ 0.05). Abbreviations: IC=Ion chromatography, Rha= Rhamnose, Ara= Arabinose, Gal= Galactose, Glc= Glucose, Xyl= Xylose, Man= Mannose, GalA = Galacturonic acid, GlcA = Glucuronic acid, TLC = Thin-layer chromatography, Ac=Acetate, DNSA= 3,5-Dinitrosalicylic acid.

To assess substrate degradation and acetate mobilization, we subjected the supernatant to acid hydrolysis followed by ion chromatography to quantify solubilized monosaccharides, and measured residual wall-bound acetate in the pellet after saponification. Treatment with *Si*GH11 alone released oligosaccharides enriched in xylose, along with minor amounts of arabinose, galactose, and galacturonic acid, consistent with cleavage of arabinoxylan side chains (Fig. 4B). Co-incubation with *Si*AXE or *Si*AXE^S226G^ did not alter the quantity or composition of the solubilized sugars, indicating that *Si*AXE does not enhance *Si*GH11 activity on insoluble xylan polymers. *Si*GH11 treatment reduced wall-bound acetate by ∼40%, consistent with the release of acetylated XOS (Fig. 4C). *Si*AXE or its inactive mutant applied individually did not reduce acetate levels, suggesting that the enzyme cannot access acetyl groups within intact xylan polymers in the cell wall matrix.

To determine whether *Si*AXE targets soluble acetylated XOS, we digested barley dAIR with *Si*GH11 and treated the resulting supernatant with water, EV control, *Si*AXE, or *Si*AXE^S226G^, followed by MALDI-ToF analysis (Fig. 4D). NaOH treatment served as a positive control for complete chemical deacetylation. While *Si*GH11 digestion followed by water or EV yielded highly acetylated XOS, subsequent treatment with *Si*AXE increased the abundance of non-acetylated species and decreased acetylated forms (Fig. 4E–F, Extended Data Fig. 5A–F). This deacetylation effect was absent in reactions with *Si*AXE^S226G^. NaOH fully deacetylated all products, confirming the expected spectral shifts.

To test whether *Si*AXE-mediated deacetylation enhances further degradation, we quantified reducing sugar release in reactions combining *Si*GH11 and *Sr*GH43 with or without *Si*AXE. Incubation with *Si*AXE significantly increased reducing sugar levels, especially when both *Si*GH11 and *Sr*GH43 were present, indicating that deacetylation facilitates complete hydrolysis of XOS (Fig. 4G–H). TLC confirmed that *Si*GH11 alone released products corresponding in mobility to Xyl2 and Xyl3, which were further digested to a product co-migrating with xylose by *Sr*GH43 (Extended Data Fig. 5G–H).

Together, these findings demonstrate that *Si*GH11 and *Si*AXE function sequentially and cooperatively to degrade monocot-derived acetylated xylan. *Si*GH11 releases acetylated XOS from the xylan backbone, and *Si*AXE deacetylates these oligosaccharides, thereby enhancing accessibility to exo-xylanases. This coordinated enzymatic activity enables complete depolymerization into non-immunogenic monosaccharides that can be efficiently utilized by *S. indica* during endophytic colonization.

### *Si*AXE controls apoplastic xylan deacetylation and supports colonization in a stage-specific manner

To investigate the biological function and secretion requirements of *Si*AXE, we generated transgenic *S. indica* strains overexpressing the full-length gene with its native signal peptide (*AXE-1* and *AXE-3*), a version lacking the signal peptide (*AXE-5*^ΔSP^), and a catalytically inactive mutant (*AXE-3*^S226G^), all driven by the *S. indica* FGB1 promoter, which is active both *in planta* and in nutrient-rich culture. Native *SiAXE* is minimally expressed in axenic culture and strongly induced during late-stage colonization of *B. distachyon* and *H. vulgare* roots (Fig. 2A–B). By contrast, transgenic *SiAXE*-OE strains showed ectopic expression in CM-grown cultures (Fig. 5A).

**Figure 5.**
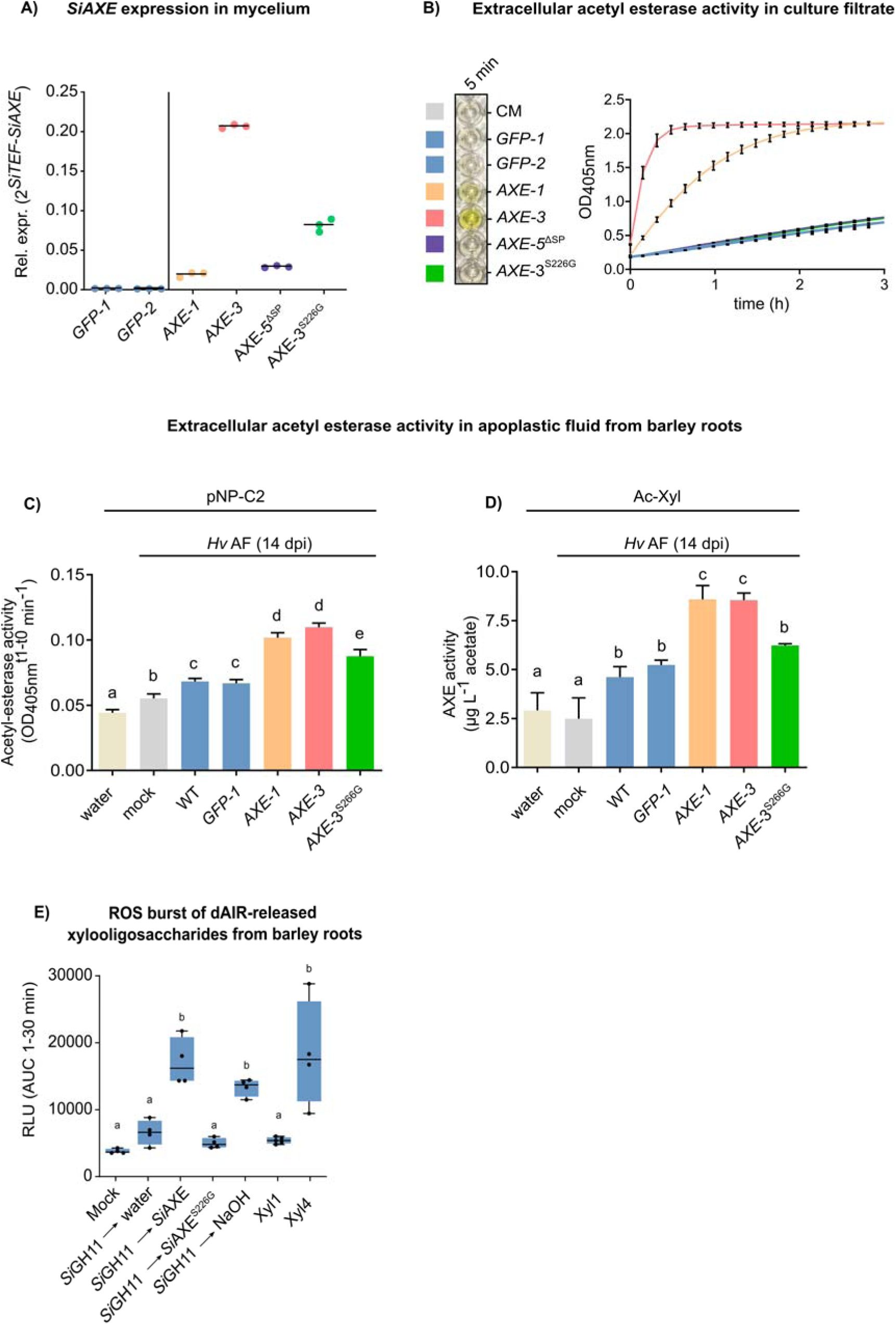
Overexpression of *SiAXE* enhances extracellular acetyl-esterase activity in axenic culture and *in planta*. **A)** qPCR validation of *SiAXE* expression in *S. indica* transformants in axenic mycelium grown in CM medium. Two independent transformants expressing full-length *SiAXE* under the control of the *S. indica* FGB1 promoter (*AXE-1*, *AXE-3*) were generated. Negative controls included *GFP*-expressing strains (*GFP-1*, *GFP-2*), a catalytic mutant (*AXE-3*^S226G^), and a transformant lacking the N-terminal signal peptide (*AXE-5*^ΔSP^). *AXE-3* displayed higher transcript levels than *AXE-1*. **B)** Acetyl-esterase activity from culture filtrates from *S. indica* transformants. *AXE-1* and *AXE-3* showed robust extracellular acetyl-esterase activity on pNP-acetate (pNP-C2), as evidenced by a color shift at OD_405_nm. No activity was detected in CM medium or filtrates from *GFP-1*, *AXE-3*^S226G^, or *AXE-5*^ΔSP^. **C–D)** Acetyl-esterase activity in apoplastic fluid (AF) isolated from barley roots at 14 dpi following inoculation with *S. indica* WT or transformants. Activity was tested using either pNP-C2 (C) or acetyl-xylan (birchwood; D). Controls included AF from mock-inoculated plants (plant baseline) and water (substrate auto-hydrolysis). Activity was measured at OD_405_nm for pNP-C2 (C), and as free acetate release from acetyl-xylan after 5 h (D). Significant differences are indicated by different letters (ANOVA + Tukey’s post hoc test, p ≤ 0.05). **F)** ROS burst assay in barley root fragments treated with soluble xylooligosaccharides generated by enzymatic digestion of barley dAIR. Supernatants of *Si*GH11 digests were harvested and further treated with *Si*AXE, controls or NaOH prior to application as depicted schematically in Figure 4D. Apoplastic ROS was measured by luminol-HRP chemiluminescence and quantified as area under the curve (AUC). Deacetylation by *Si*AXE and NaOH significantly increased ROS accumulation, unlike water or *Si*AXE^S226G^. Mock-treated roots and those treated with xylose (Xyl1) or xylotetraose (Xyl4) served as controls (see Fig. S9 for kinetics).

*AXE-1* and *AXE-3* secreted functional *Si*AXE protein into the medium, as confirmed by robust acetyl esterase activity in culture filtrates that correlated with transcript levels. No such activity was detected in control strains expressing *GFP* (*GFP-1*, *GFP-2*), the catalytically inactive *AXE-3*^S226G^, or the secretion-deficient *AXE-5*^ΔSP^ (Fig. 5B). Western blot analysis confirmed the presence of recombinant *Si*AXE-HA in both the culture supernatant and mycelium of OE strains, while removal of the signal peptide abolished extracellular activity and led to intracellular accumulation of a smaller, non-glycosylated *Si*AXE form, consistent with mislocalization and secretion failure (Extended Data Fig. 6A-C). Enzymatic deglycosylation of mycelial protein from *AXE-1* and *AXE-3* led to the appearance of a smaller band, comparable in size to that observed in *AXE-5*^ΔSP^, indicating that native *Si*AXE is glycosylated. To assess *in planta* activity, we isolated apoplastic fluid (AF) from barley roots that were either mock-inoculated or colonized with wild-type (WT) *S. indica* or the different fungal transformants. AF from *S. indica* WT and *GFP-1*–inoculated roots showed increased acetyl-esterase and acetyl-xylan esterase activity compared to the mock-inoculated control. This activity was markedly elevated in roots colonized by *AXE-1* and *AXE-3* but remained low in *AXE-3*^S226G^ (Fig. 5C–D), demonstrating that *Si*AXE contributes to apoplastic acetyl-xylan ester hydrolysis during colonization and that its activity can be modulated through transgenic expression.

To evaluate *Si*AXE function *in planta*, we measured apoplastic ROS production in barley roots as a functional readout for the presence of immunogenic deacetylated XOS. Supernatants from *Si*GH11-digested barley dAIR triggered a modest ROS response, which was significantly amplified after enzymatic deacetylation by *Si*AXE or chemical deacetylation with NaOH (Fig. 5E, Extended Data Fig. 7). Consistently, application of defined oligosaccharides such as xylotetraose (Xyl4) induced ROS production, whereas the monosaccharide xylose (Xyl1) did not. Size-exclusion chromatography of hydrolysates further showed that deacetylated XOS of various lengths elicited stronger ROS responses (Extended Data Fig. 8), providing indirect but functional *in planta* evidence of *Si*AXE activity.

To evaluate the role of *Si*AXE in fungal colonization, we additionally generated a *Si*AXE knockout (KO) strain (*axe-1*) via homologous recombination (Extended Data Fig. 9). For this, we established a homokaryotic (hk) *S. indica* strain (hk#58 WT), which contains only a single nucleus type per cell, ensuring genetic uniformity and facilitating stable gene targeting. We then compared *S. indica* OE and KO strains with their respective dikaryotic and homokaryotic control strains across a time course of barley root infection. At 3 dpi, *SiAXE* overexpression led to increased intraradical colonization, as indicated by elevated *SiTEF* transcript levels (Fig. 6A). By 7 and 14 dpi, colonization levels converged across all transformants, suggesting that early *SiAXE* OE promotes initial intracellular progression, while endogenous expression may suffice at later stages. Expression of the barley defense marker *HvPR10* was also transiently elevated at 3 dpi in roots colonized by *SiAXE*-OE transformants, indicating a mild immune response concurrent with enhanced fungal proliferation (Fig. 6C). This suggests that premature *SiAXE* expression may enhance acetate availability through oligosaccharide deacetylation in the apoplast, thereby supporting early nutrient acquisition while transiently activating host immunity. In contrast, analysis of the *SiAXE* KO strain (*axe-1*) in the homokaryotic background, in which *SiAXE* expression was undetectable *in planta*, revealed normal colonization levels at 3 and 7 dpi but significantly reduced fungal intracellular proliferation at 14 dpi compared to the homokaryotic control strain (Fig. 6D–E). Together, these findings indicate that *Si*AXE promotes early-stage colonization when overexpressed and is required for sustained fungal growth during late-stage colonization (Fig. 7).

**Figure 6.**
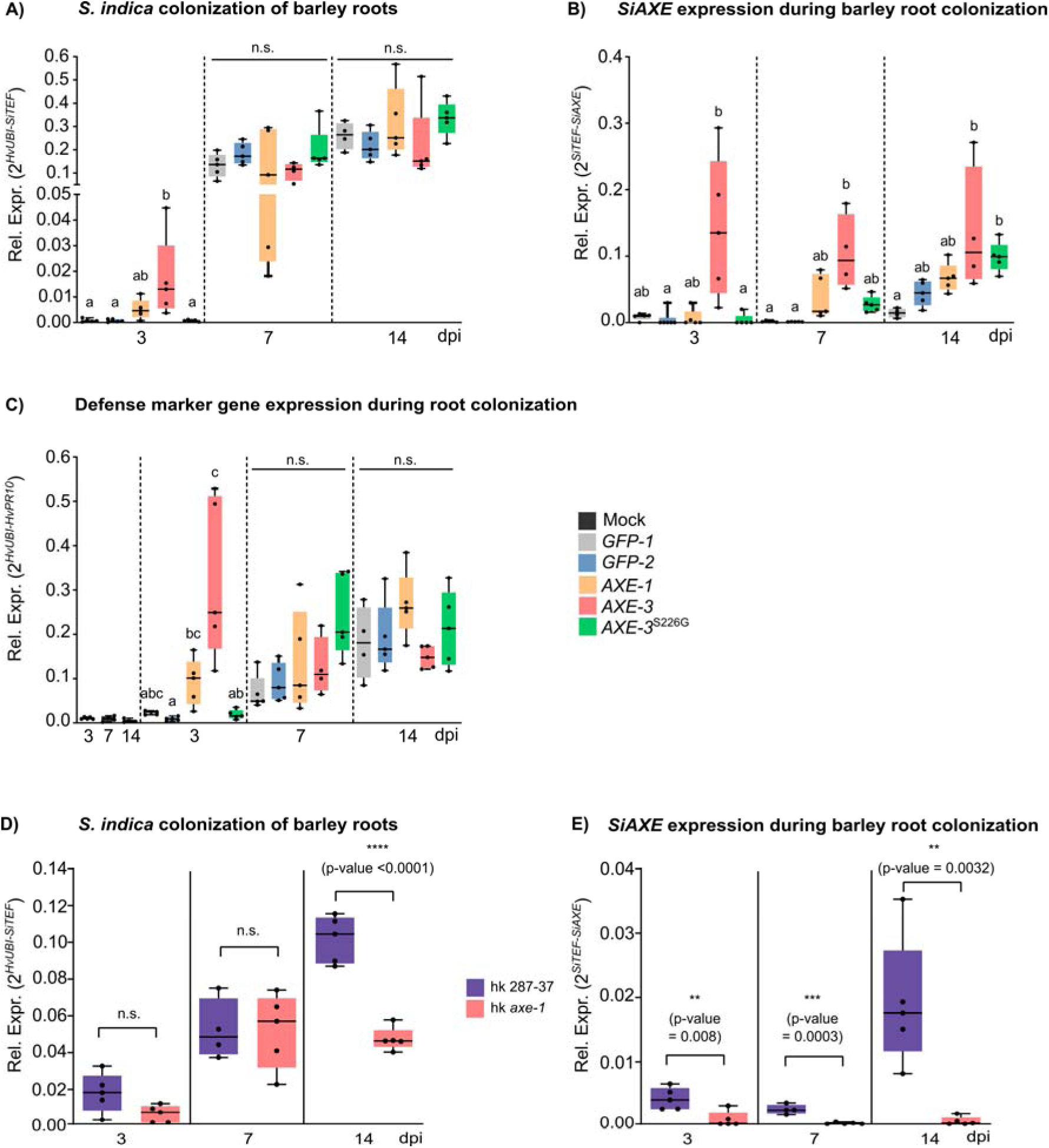
Overexpression of *SiAXE* enhances early root colonization, while loss of function reduces colonization at later stages. **A–C)** Barley roots were inoculated with *S. indica* transformants expressing full-length *SiAXE* (*AXE-1*, *AXE-3*), a catalytic mutant (*AXE-3*^S226G^), or cytosolic GFP (*GFP-1*, *GFP-2*) as controls. Colonization was monitored at 3, 7, and 14 days post-inoculation (dpi). **A)** Quantification of root colonization via relative expression of the fungal marker gene *SiTEF* normalized to the barley reference gene *HvUBI*. Colonization by *AXE-3* is significantly elevated at 3 dpi compared to controls, but differences are not observed at 7 or 14 dpi. **B)** *SiAXE* expression levels *in planta*, showing early induction in *AXE-3* at 3 dpi, and in both *AXE-1* and *AXE-3* at 7 dpi. At 14 dpi, all *S. indica* strains exhibit *SiAXE* expression, consistent with its induction during the late saprotrophic stage. **C)** Expression of barley defense marker *HvPR10*. Induction is observed at 3 dpi in plants colonized by *AXE-1* and *AXE-3*, with no significant differences at later time points. Different letters in panels A–C indicate statistically significant differences (ANOVA + Tukey’s post hoc test, p ≤ 0.05). **D–E)** Barley inoculated with *S. indica* deletion mutant *axe-1* (deficient in *SiAXE*) and its genetic background control (hk K287-37) containing a wild-type copy of *SiAXE*. **D)** Colonization levels measured by *SiTEF/HvUBI* ratio. *axe-1* exhibits significantly reduced colonization at 14 dpi compared to the control. **E)** *SiAXE* expression is strongly reduced in *axe-1* across all time points, while hk K287-37 shows normal temporal induction, peaking at 14 dpi. Asterisks indicate significant differences (two-tailed Student’s *t*-test): n.s. = not significant; *p* _≤_ *0.01 (**), p* _≤_ *0.001 (\*\*\**).

**Figure 7.**
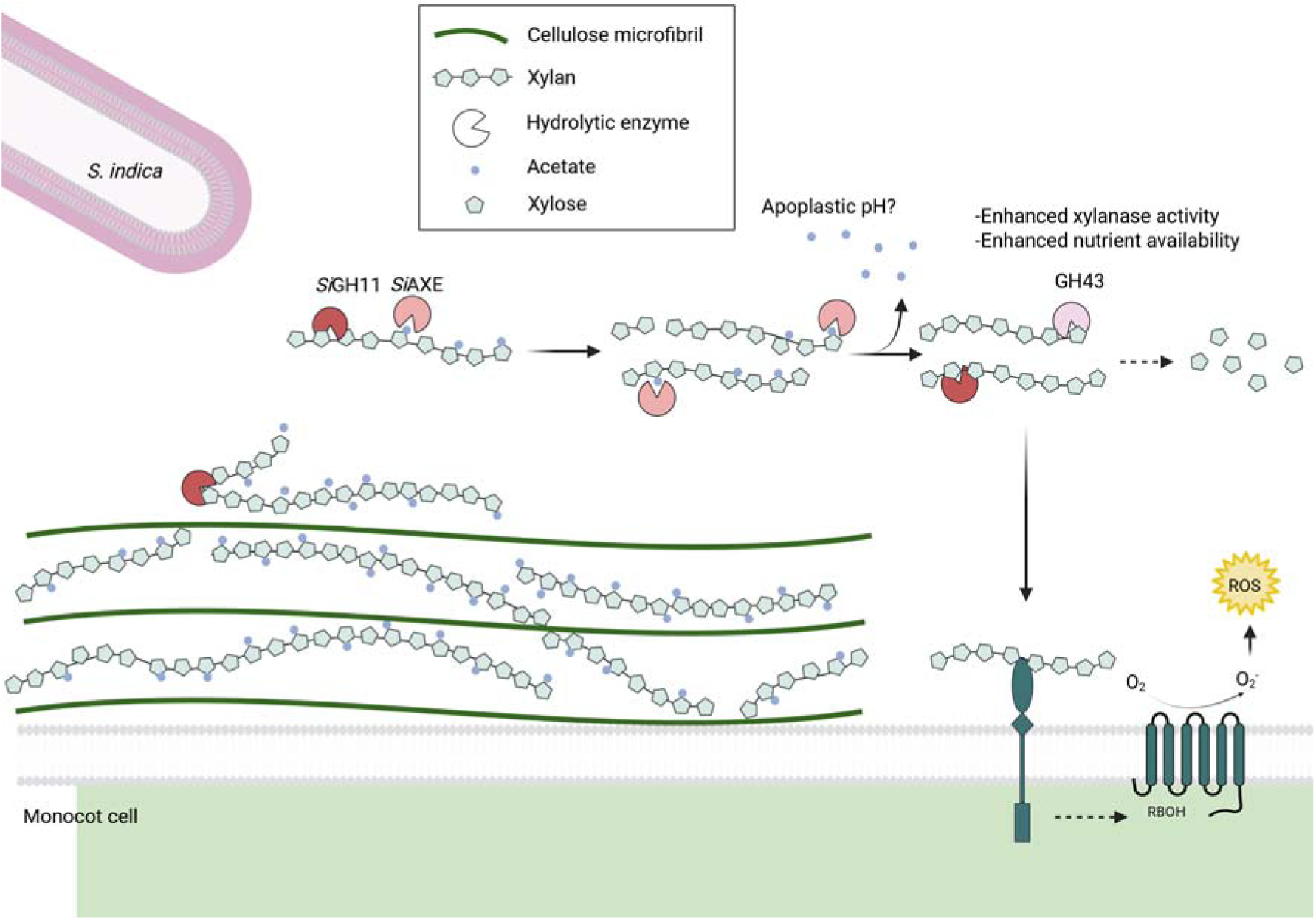
Model of cooperative activity between *Si*AXE and *Si*GH11 in the enzymatic digestion of acetyl-xylan from monocot cell walls. During the late saprotrophic stage of root colonization, *S. indica* secretes a cocktail of cell wall–modifying enzymes (CAZymes) into the apoplast. These include co-regulated members of WCNA module 7, notably the endo-xylanase *Si*GH11 and the XynE-like acetyl-xylan esterase *Si*AXE. Together, these enzymes degrade the acetylated xylan backbone of monocot cell walls by releasing and deacetylating xylooligosaccharides (XOS). XOS of varying lengths can act as damage-associated molecular patterns (DAMPs), triggering rapid production of apoplastic reactive oxygen species (ROS)—a hallmark of pattern-triggered immunity (PTI). Deacetylated XOS are particularly potent inducers of ROS. *Si*AXE hydrolyzes acetyl groups from XOS, releasing free acetate into the apoplast. This may modulate apoplastic pH and provide acetate as an auxiliary carbon source for the fungus. Critically, deacetylation of XOS by *Si*AXE renders them susceptible to complete hydrolysis by a combination of endo-xylanases (e.g., *Si*GH11) and exo-xylanases (e.g., GH43 family), generating non-immunogenic xylose for fungal nutrition. This host-adapted enzymatic strategy facilitates immune-compatible degradation of host hemicellulose and supports mutualistic colonization of monocot roots.

## Discussion

Successful intracellular colonization of plant roots by mutualistic fungi requires tightly regulated, host-adapted, and stage-specific remodeling of host cell walls. Our study identifies a transcriptionally coordinated enzymatic program in *S. indica* that underlies this adaptation during colonization of monocot hosts. Specifically, we demonstrate that the glycoside hydrolase *Si*GH11 and the acetyl-xylan esterase *Si*AXE act cooperatively to degrade *O*-acetylated xylan, a major component of barley root cell walls. These enzymes are co-expressed as part of a late-stage, monocot-specific gene module (Module 7; Fig. 1 and Fig. 2), which is enriched in secreted CAZymes and sugar transporters. This module reflects a broader metabolic transition associated with localized wall remodeling and host-derived carbon acquisition during intracellular accommodation. Consistent with its transcriptional profile, *SiGH11* was previously identified as a marker of the cell death–associated phase of barley colonization, during which *S. indica* primarily occupies senescing or dying host cells in a manner reminiscent of saprotrophic growthL^22^.

Biochemical analysis confirms that *Si*GH11 functions as a canonical endo-1,4-beta-xylanase, releasing soluble acetylated XOS, ultimately xylobiose and xylotriose, from native barley xylan. Mass spectrometry analysis showed that *Si*AXE efficiently deacetylates these *Si*GH11-derived XOS, generating fragmentation patterns comparable to those produced by chemical deacetylation. Notably, acetate levels remained unchanged in barley root cell walls treated with *Si*AXE alone, indicating that its activity is restricted to soluble rather than wall-bound substrates. These findings support a model in which *Si*GH11 and *Si*AXE act sequentially to degrade acetylated xylan in the apoplast. This two-step mechanism appears to operate *in planta*. Deacetylation of native *Si*GH11-derived XOS enhanced apoplastic reactive oxygen species accumulation in barley roots, which is consistent with increased exposure of damage-associated molecular pattern motifs ^43,44^. Co-incubation of barley cell walls with *Si*GH11 and the exo-xylanase *Sr*GH43, which digests XOS from the non-reducing end, resulted in the accumulation of non-immunogenic xylose as the terminal product. Notably, sugar release by *Si*GH11 and *Sr*GH43 was enhanced by enzymatic deacetylation with *Si*AXE, supporting the notion that acetyl groups sterically hinder glycoside hydrolase activity ^38,41^. Although *Sr*GH43 is of bacterial origin, several GH43 family enzymes are encoded within the same late-stage *S. indica* gene module. This suggests that a similar enzymatic cascade may function *in planta* to complete xylan depolymerization and minimize immune activation.

*Si*GH11 and *Si*AXE function within a broader carbohydrate-degrading network encoded by module 7, which includes multiple endo-xylanases (GH10 and GH11), acetyl-xylan esterases (CE1 and CE3), α-L-arabinofuranosidases (GH43_5 and GH62), an exo-xylanase (GH43_36), and a (methyl)glucuronidase (GH115) (Fig. 7, Extended Data Fig. 10). Although some GH10 and GH11 enzymes have been reported to exhibit exo-xylanase activity, such activity appears limited in *Si*GH11 (PIIN_05889). Our biochemical assays and ROS burst experiments indicate that *Si*GH11 can partially degrade Xyl3, consistent with low exo-β-1,4-xylosidase activity, but not sufficient to fully depolymerize XOS (Extended Data Fig. 5G–H). This suggests that complete conversion of XOS to monosaccharides *in planta* likely requires cooperation with dedicated exo-acting enzymes such as GH43_36, which is co-expressed in the same transcriptional module.

Together, these enzymes likely act synergistically to deconstruct the structurally complex acetylated xylan typical of monocot cell walls, releasing xylose, arabinose, and acetate as primary breakdown products. Co-expressed sugar transporters, including PIIN_03369 and PIIN_11722 (homologous to *Xylhp*, *XLT1*, *CDT-1*, and *CDT-2*) are likely involved in the uptake of the released mono- and oligosaccharides ^45–47^. Module 7 also encodes key components of the pentose catabolic pathway (PCP), such as a D-xylose reductase (PIIN_02595) and two alcohol dehydrogenases (PIIN_05868 and PIIN_05869), which may enable *S. indica* to funnel imported pentoses into the pentose phosphate pathway. This organization resembles metabolic strategies described in *Aspergillus niger* and *Magnaporthe oryzae* ^48–51^ (Extended Data Fig. 10). Despite the immunogenic potential of deacetylated XOS, *S. indica* triggers only weak immune responses during colonization. This may reflect a concerted strategy in which secreted glycoside hydrolases and esterases rapidly fragment xylan, while co-expressed sugar transporters sequester monosaccharides and immune-active oligosaccharides before they accumulate in the apoplast (Fig. 7).

The functional relevance of this tightly regulated enzymatic module is further supported by gain- and loss-of-function analyses. Overexpression of *SiAXE* enhances early-stage root colonization and is associated with a mild immune response, whereas deletion of the gene impairs fungal proliferation during the later stages of intracellular colonization. These findings point to a dual role for *Si*AXE in host cell wall remodeling and metabolic integration during endophytic growth. More broadly, our data show that CAZyme deployment occurs within a monocot-specific transcriptional program that also encompasses secreted effectors and nutrient transporters. This coordinated regulation highlights how mutualistic fungi like *S. indica* integrate structural, immunological, and metabolic functions to establish and maintain compatibility within specific plant lineages.

In summary, our work reveals a host- and stage-specific enzymatic strategy by which a mutualistic fungus accesses carbon from complex, acetylated xylan in monocot cell walls and facilitates endophytic colonization by locally and selectively weakening structural barriers. We propose that concerted enzymatic activity by members of module 7 supports colonization by specifically targeting *O*-acetylated xylan in barley root walls. This degradation ultimately yields pentoses and acetate as nutrient sources, while concurrently limiting ROS production and reducing the immunogenic potential of intermediate XOS. These hydrolysis products alter the biochemical environment of the apoplast, dampening immune responses and promoting fungal establishment. The coordinated expression and functional specialization of *Si*GH11 and *S*iAXE, a highly specific acetyl-esterase that releases acetate from soluble acetylated XOS, alongside exo-xylanases, sugar transporters, and metabolic enzymes, exemplifies how modular enzyme systems are co-opted and fine-tuned across the saprotrophy-to-symbiosis continuum to remodel host walls, integrate nutrient metabolism, and enable colonization within the immunologically sensitive apoplastic niche.

## Methods

### Plant material and growth conditions

Seeds of *Hordeum vulgare* cv Golden Promise were surface sterilized with 6% sodium-hypochlorite for 1 h and washed six times for 30 min with sterile milliQ-water (MQ-H_2_O). Seed coats were removed gently and seeds were germinated on wet filter paper for three days in the dark at 21 °C. For fungal colonization, germinated seedlings were transferred to sterile WECK jars containing 100 ml 1/10 PNM (1 ml L^-1^ 500 mM KNO_3_, 1 ml L^-1^ 367 mM KH_2_PO_4_, 1 ml L^-1^ 144 mM K_2_HPO_4_, 1 ml L^-1^ 2 M MgSO_4_, 1 ml L^-1^ 200 mM Ca(NO_3_)_2_ 4H_2_O, 1 ml L^-1^ 428 mM NaCl, 5 ml L^-1^ 10 mM Fe-EDTA (Fe^2+^), pH 5.6, 10 ml L^-1^ 1 M MES, 1.2 % (w/v) Gelrite) and transferred to light under long-day conditions (day/night cycle of 16/8 h, 22 °C/18 °C, light intensity of 108 μmol/m^2^*s) for one day and then inoculated with *S. indica* chlamydospores and continued to be grown for 3-, 7- and 14 dpi prior to washing and harvesting of roots. For ROS burst assays, seeds were sterilized and germinated on filter paper and grown for four days prior to root preparation.

### Cultivation of *S. indica*, chlamydospore isolation and barley inoculation

*S. indica* WT (strain DSM11827) and transformants were grown in CM medium (20 g L^-1^ Glucose, 2 g L^-1^ peptone, 1 g L^-1^ yeast extract, 1 g L^-1^ caseine hydrolysate, 1 ml L^-1^ microelements: 6g MnCl_2_ 4H_2_O, 1.5 g H_3_BO_3_, 2.65 g ZnSO_4_ 7H_2_O, 750 mg Kl, 2.4 mg Na_2_MO_4_ 2H_2_O, 130 mg CuSO_4_ 5H_2_O-, 50 ml L^-1^ salt solution: 120 g NaNO_3_, 10.4 g KCl, 10.4 g MgSO_4_ 7H_2_O, 30.4 g KH_2_PO_4_ and 15 g L^-1^ agar in the case of solid medium), containing hygromycine (80 µg ml^-1^) in the case of transformed *S. indica*. *S. indica* transformants were maintained by transferring a plug (5 mm diameter) stamped out from an CM agar plate containing actively growing mycelium to a fresh plate and further grown for four weeks at 28°C. Chlamydospore isolation was done as previously described ^32^. Briefly, 5 ml tween water (0.002 % v/v) was added to a four-week old plate and gently scraped with a scalpel and the spore suspension filtered through a miracloth funnel (22-25 µm pore size) and collected in a 50 ml falcon tube, centrifuged for 7 min at 3350 g using a swing-out rotor, the supernatant decanted and the pellet washed twice with 10 ml tween water and finally resuspended in 10 ml MQ-H_2_O and spore concentration counted using a Neubauer improved counting chamber and diluted to the desired concentration. For plant inoculation, germinated barley seedlings were prepared and inoculated with 3 ml of either sterile water as mock control or *S. indica* chlamydospores (500.000 spores ml^-1^). Roots were harvested at 3-, 7- and 14 dpi, washed thoroughly in ice-cold water to remove extraradical fungal hyphae, briefly dried on tissue paper and frozen in liquid N_2_ for protein or RNA extraction. Four barley plants were used per jar and pooled per biological replicate.

### Isolation of *S. indica* mycelium and culture filtrate

For isolation of rapidly growing, metabolically active mycelium, a starter culture of 50 ml CM medium in a 200 ml Erlenmeyer flask was inoculated with chlamydospores from a four-week old plate and grown for one week at 120 rpm at 28°C. The growing mycelium was harvest by pouring the culture through a miracloth funnel, which retains the mycelium. This was subsequently surface washed with 50 ml 0.9 % NaCl and MQ-H_2_O and then homogenized in a blending mixer containing 20 ml fresh CM medium. The homogenized mycelium was transferred to 180 ml CM medium in a 500 ml Erlenmeyer flask and grown for 2 days at 28°C. Then, mycelium was harvested again and the culture filtrate (CF) put aside for enzymatic assays and immunodetection by western blot. The remaining mycelium was surface-washed with 50 ml 0.9% NaCl and 50 ml MQ-H_2_O, dried briefly on tissue paper and frozen in liquid N_2_ for protein or RNA extraction.

### Apoplastic fluid isolation of barley roots

Apoplastic fluid (AF) was isolated from barley seedlings as described previously ^52^. Germinated barley seedlings were inoculated with *S. indica* on 1/10 PNM medium and per replicate, 40 barley seedlings were harvested at 14 dpi, which yielded approximately ∼0.5 ml AF per replicate. Harvested roots were surface washed thoroughly in ice-cold water to remove external fungal material, cut into pieces of 2 cm, transferred to ice cold water and vacuum infiltrated (five cycles of15 min 250 mbar, 1.5 min ATM). Then, the roots were gently dried on tissue paper and transferred into a 20 ml syringe in a 50 ml falcon tube, centrifuged for 15 min at 4 °C, 2000 rpm (lowest de- and acceleration) to collect AF in the falcon tube. The isolated AF was stored at −20°C or immediately for enzymatic assays.

### qRT-PCR

For qRT-PCR, mock-inoculated or *S. indica*-colonized barley roots and axenic *S. indica* mycelium were ground to a fine powder an 200 mg used for RNA isolation and cDNA synthesis as described previously ^53^.

### Recombinant protein production in *E. coli*

For heterologous expression of *S. indica* genes, the coding sequences of *SiAXE* (PIIN_07653), *SiPLL* (PIIN_06162) and *SiGH11* (PIIN_05889) were amplified without N-terminal signal peptide from cDNA prepared from colonized barley roots and cloned via Gibson assembly (NEB; #E2611) into QIAexpress pQE-80L expression vector (Qiagen). As control, a catalytic mutant of *SiAXE* was generated replacing Serine226 with Glycine (S226G) using the Q5 Site-Directed Mutagenesis Kit (NEB; #E0554S). Recombinant protein was isolated from transformed *E. coli* BL21(DE3)pLysS (Promega) via their N-terminal ^6x^His-tag by Ni-NTA agarose resin using imidazole as described previously ^54^ and subsequently dialyzed against protein dialysis buffer (50 mM K_2_HPO_4_-KH_2_PO_4_, pH=6.9), protein amount quantified with Bradford reagent (Bio-Rad) and diluted to 50 µg ml^-1^ with dialysis buffer for *in vitro* enzyme assays. As additional control, the same steps were performed with a culture containing the empty vector (EV). Presence and purity of recombinant protein was confirmed by immunoblotting after SDS-PAGE using KPL HisDetector Nickel-HRP (Sera Care; #5820-0001).

### Overexpression of *SiAXE* in *S. indica*

The coding sequence of *SiAXE* was amplified via PCR from cDNA prepared from colonized barley roots either full length or without N-terminal signal peptide. A catalytic mutant (S226G) of *SiAXE* was generated using the Q5 Site-Directed Mutagenesis Kit (NEB; #E0554S). The PCR products were cloned via Gibson assembly (NEB; #E2611) into the K100 expression vector ^32^, expressing –C-terminal HA-tagged *SiAXE* under the control of the *SiFGB1* promotor (see table S2 for information on primer sequences and plasmids). Transformation of *S. indica* was done as previously described ^32^. Per transformation, 10 µg of blunt-end linearized plasmid was purified from agarose-gel and used for PEG-mediated transformation of *S. indica* protoplasts at a concentration of 10^7^-10^9^ mL^-1^ protoplasts obtained from dikaryotic (dk) or homokaryotic (hk#58 WT) *S. indica*. Individual, regenerated *S. indica* transformants were transferred one by one to fresh CM + hygromycine (80 µg ml^-1^) agar after 7-21 days and kept until further use. The individual *S. indica* transformants were transferred to new CM +Hyg agar plates every 4-6 weeks. For DNA extraction, a 1 cm^2^ section of growing mycelium was removed with a scalpel from the surface of the plate and used for extraction of genomic DNA to determine the genotype (dk or hk) by testing via PCR for the presence of the multiallelic homeodomain (HD) encoding genes PIIN_09915 (HD1.1), PIIN_ 09916 (HD2.1), PIIN_09977 (HD1.2) and PIIN_09978 (HD2.2) by which they can be distinguished as dikaryotic or homokaryotic ^55^. Per transformation, five individual strains were tested for expression of *SiAXE* and extracellular acetyl-esterase activity in axenic culture prior to further use. See table S3 for transgenic *S. indica* transformants.

### Generation of mutant *axe-1* in homokaryotic *S. indica* via homologous recombination

The generation of the *axe-1* null mutant is described in detail in Figure S11. The *axe-1* mutant was created via homologous recombination using *S. indica* genomic border sequences of 2000 bp flanking the *SiAXE* genomic locus, termed left border (LB) and right border (RB). These were amplified from *S. indica* genomic DNA and cloned up and downstream of a hygromycine resistance cassette (HygR) comprising the TEF promotor from *S. vermifera* (SvTEFPro), the hygromycin resistance gene and the Nos terminator (NosT), into plasmid K287 (see table S2 for plasmids). Prior to transformation, plasmid K287 was digested into LB and RB and a ∼ 600 bp overhang for homologous recombination using *Pme*I+*Sac*II and *AsiS*I, respectively and the fragments gel-purified after agarose gel electrophoresis. Per border, 10 µg linear DNA was used for PEG-mediated transformation of protoplasts obtained from homokaryotic (hk#58 WT) *S. indica* as described previously ^32^. To verify that the LB-HygR-RB cassette replaced the *SiAXE* genomic locus, genomic DNA was isolated from all regenerated colonies from the transformation reactions and tested using various PCR reactions (Figure S11). A mutant (K287-13; *axe-1*) and a genetic background control (K287-37) were selected and RNA was extracted from axenic mycelium and absence of *SiAXE* expression confirmed.

### Extraction of protein from *S. indica* mycelium or colonized barley roots

For protein extraction, *S. indica* mycelium or colonized roots were ground with mortar and pestle under liquid N2 to a fine powder and 200 mg FW transferred per sample to a 2 ml tube. Then, 500 µl of pre-cooled TKMES extraction buffer (100 mM tricine; pH=7.5, 10 mM KCL, 10 mM MgCl2, 1 mM EDTA, 10% w/v sucrose, freshly added 0.2 % v/v Triton X-100, 1 mM DTT, 100 µg ml^-1^ PMSF, 1x protease inhibitor cocktail (Sigma Aldrich; #P9599) was added and vortexed vigorously for 20 seconds. For samples that were used for xylanase assays, sucrose was omitted from the buffer (TKME). The homogenate was centrifuged at 10000 g for 10 minutes at 4°C to pellet cell debris and the supernatant harvested, protein amount quantified with Bradford reagent (Bio-Rad) and diluted to 0.3 mg ml^-1^ with the appropriate buffer for *in vitro* enzyme assays. Presence of *Si*AXE-HA was confirmed in mycelium protein or culture filtrate after SDS-PAGE and immunoblotting using anti-HA antibody (Sigma; #H9658-2ML) as primary and anti mouse HRP (Sigma; #A2304) as secondary antibody.

### Deglycosylation of *S. indica* mycelium protein

*S. indica* mycelium protein extracts from transformants overexpressing *SiAXE* (*AXE-1*; *AXE-3*) and the truncated version without N-terminal signal peptide (*AXE-5*^ΔSP^) were enzymatically deglycosylated using denaturing conditions with the Protein Deglycosylation Mix II (NEB; #P6044S). A total of 20 µg mycelium protein was digested in a 50 µl reaction containing 5 µl Deglycosylation Mix II and 5 µl buffer 2, incubated at 25°C for 30 min and then at 37°C for 16h, prior to analysis by SDS-PAGE and immunoblotting.

### *In vitro* esterase and lipase activity assay

Esterase and lipase activity was determined in protein extract from axenic *S. indica* mycelium and CF, AF from mock-inoculated or *S. indica*-colonized roots or with the recombinant protein *SiAXE* and *Si*PLL. As substrates, *para*-nitrophenyl (pNP) carboxyl esters of following carbon chain length were used: acetate (C2; Merck; #46021), butyrate (C4; Merck; #N9876), hexanoate (C6; Th. Geyer; #BD244418), octanoate (C8; Merck; #S574120), decanoate (C10; Merck, #N0252), dodecanoate (C12; Merck; #61716), myristate (C14; Merck; #70124) and palmitate (C16; Merck; #N2752). Substrates were prepared as 1 mg ml^-1^ stocks in ethanol from serial dilutions and heated to 70 °C before pipetting to ensure solubility. Recombinant *Si*PLL and commercial *R. oryzae* lipase (*Ro*LIP) (Merck; #62305) served as control for lipase activity. Reactions were set up with 90 µl reaction mix (10% (v/v) substrate (1 mg ml^-1^, in ethanol), 1 % (v/v) Triton X-100, in Tris-HCL, 25 mM, pH=7.5) and 10 µl protein, protein extract or CF. For AF, 150 µl reaction mix and 50 µl AF were used. 100 µl of every reaction was transferred to a transparent 96-well plate (Greiner), the plate sealed and shift in absorbance measured regularly at OD 405 nm in a TECAN SPARK 10M microplate reader over 20 h. Control reactions of non-enzymatic hydrolysis were set up with appropriate buffers, respectively. The enzyme-substrate combinations that were tested are indicated in the legends and the respective figures. Per reaction conducted together, similar amounts of protein, protein extracts and substrate were used to ensure comparability.

### Cell-wall polymers and oligosaccharides

The following CW polymers or oligosaccharides were used. Birchwood acetyl-Xylan (Megazyme; #P-ACXYL), beechwood xylan (Megazyme; #P-XYLNBE), shrimp shell chitin (Merck; #C7170), cellulose (Merck; #C6414), glucose-pentaacetate (Ac-Gluc; Merck; #285943), galactose-pentaacetate (Ac-Gal; Merck; #134031), xylose (Xyl1; Roth; #5537.1), xylobiose (Xyl2; Megazyme; #O-XBI), xylotriose (Xyl3; Megazyme; #O-XTR), xylotetraose (Xyl4; Megazyme; #O-XTE), xylopentaose (Xyl5; Megazyme; #O-XPE), xylohexaose (Xyl6; Megazyme; #O-XHE).

### *In vitro* acetyl-xylan esterase activity assays

Acetyl-xylan esterase (AXE) activity was determined in protein extracts or AF from mock-inoculated and colonized barley roots, or with recombinant *SiAXE* and controls. Activity was tested with birchwood acetyl-Xylan. For root protein extracts, reactions contained 170 µl substrate (10 mg ml^-1^, in Tris-HCL, 25 mM, pH=7.5) and 30 µl protein extracts. For AF, reactions contained 2.5 µl substrate (100 mg ml^-1^, in Tris-HCL, 25 mM, pH=7.5), 22.5 µl Tris-HCL (25 mM, pH=7.5) and 25 µl AF. The reactions were incubated at 28°C under continuous shaking for 16h. For recombinant *SiAXE*, activity was also tested with Ac-Gluc and Ac-Gal in a reaction containing 135 µl Tris-HCL (25 mM, pH=7.5), 7.5 µl substrate (100 mg ml^-1^, in Tris-HCL, 25 mM, pH=7.5) and 7.5 µl protein and incubated at 28°C under continuous shaking for 1h. After incubation, the samples were heat inactivated for 5 min at 95°C, centrifuged for 5 min at 13.000 rpm and 5-70 µl of the supernatant used to quantify released acetate using the Acetic Acid Assay Kit (Megazyme; #K-ACET) in a 100 µl reaction adapted to a 96-well plate format according to the supplier’s instructions. Control reactions of non-enzymatic hydrolysis were set up with appropriate buffers, respectively. The enzyme-substrate combinations that were tested are indicated in the legends and the respective figures. Per reaction conducted together, similar amounts of protein, protein extracts and substrate were used to ensure comparability.

### *In vitro* xylanase activity assays

Xylanase activity was determined either in protein extracts from mock-inoculated and colonized barley roots, or with recombinant *Si*GH11 and *Sr*GH43 β-D-xylosidase (exo-xylanase, Megazyme; #E-BXSR-1KU). For xylanase activity in roots, 100 µl protein extract (0.3 mg ml^-1^, in TKME buffer), was incubated with 50 µl beechwood xylan (10 mg ml^-1^). As substrates for recombinant enzymes, birchwood acetyl-xylan, beechwood xylan, konjak glucomannan, shrimp shell chitin, cellulose or a mixture of the linear XOS, Xyl1-Xyl6 was used. Reactions with CW polymers were set up with 100 µl substrate (1-10 mg ml^-1^, in water), 25 µl Tris-HCL (25 mM, pH=7.5), each 25 µl *Si*GH11, *Sr*GH43 or EV in a total volume of 175 µl. For XOS, a 30 µl equimolar (5 mM) mixture in Tris-HCL (25 mM, pH=7.5) was mixed with each 15 µl *Si*GH11, *Sr*GH43 or EV in a total volume of 60 µl. All enzyme digests were incubated at 28°C under continuous shaking for 16 h and terminated at 95°C for 5 min, centrifuged at 13000 rpm for 5 min and the supernatants used for DNSA quantification of reducing end sugars, or thin layer chromatography (TLC) analysis of hydrolysates. Control reactions of non-enzymatic hydrolysis were set up with appropriate buffers, respectively. The enzyme-substrate combinations that were tested are indicated in the legends and the respective figures. Per reaction conducted together, similar amounts of protein, protein extracts and substrate were used to ensure comparability.

### Combined *in vitro* enzyme assay of *Si*GH11, *Sr*GH43, *Si*AXE

The combined enzyme reactions contained 100 µl substrate (10 mg ml^-1^ barley dAIR or beechwood xylan), each 10 µl combined recombinant enzymes or dialysis buffer in a total volume of 130 µl and incubated under continuous shaking at 28°C for 1.5h, 5h and 17h. Then, reactions were heated at 95°C for 5 min, centrifuged at 13000 rpm for 5 min and the supernatant used for quantification of reducing end sugars with the DNSA method or for thin-layer chromatography analysis.

### Thin-layer chromatography

The supernatants of enzymatic hydrolysis of CW polymers or XOS were applied to thin-layer chromatography (TLC) to analyze the soluble oligosaccharide products. Samples were applied 1 cm from the bottom of a TLC Silica gel 60 F254 plate (Merck; #1.05554.0001) and per sample 2-10 µl were applied and allowed to air dry in between application steps. The plates were developed in a closed glass tank with 500 ml running buffer (208 ml n-buthanol, 166 ml methanol, 83.2 ml 28% ammonium hydroxide, 41.6 ml water) for 30-45 min. The plates were air dried and briefly placed in developer solution (80 ml acetone, 12 ml 85% phosphoric acid, 1.6 g diphenylamine, 1.6 g aniline), air dried again and placed in an oven at 80°C for 20-30 min.

### Measurement of reducing-end sugars

For determination reducing sugars after xylanase assays, the dinitrosalicylic acid (DNSA) method for quantification of colorimetric complexes formed by reducing sugars with 3,5-dinitrosalicylic acid was used ^56^. One volume of sample was mixed with 1.5 volume (v/v) of the DNSA color reagent and incubated for 10 min at 95°C, cooled on ice and absorbance determined at OD 540 nm.

### Endo-xylanase activity assay

Endo-xylanase activity was tested for recombinant *Si*GH11 and *Aspergillus niger* hemicellulase C (AnHemC; Merck, #H2125) as positive control. Assays were performed using birchwood azo-xylan (Megazyme; #S-AXBP) according to supplier’s instruction. Briefly, 500 µl substrate (10 mg ml^-1^), 400 µl phosphate buffer (10 mM, pH=6.0) and 100 µl enzyme were mixed and incubated for 0.5 h or 16 h. Absorbance shift, indicative of endo-xylanase activity, was determined in aliquots of 200 µl. To this end, 500 µl (95% v/v) ethanol was added, vortexed and incubated at 21 °C for 5 min, centrifuged at 3000 rpm for 10 min and OD 590 nm recorded in 100 µl supernatant.

### Isolation of cell wall material from barley roots

Mock-inoculated barley roots were harvested 14 dpi and lyophilized and milled to a fine powder using a Retsch MM400 mixer mill. Then, alcohol insoluble residue (AIR) was prepared as previously described ^57^. AIR was delignified with peracetic acid based as previously described ^58^. Briefly, AIR samples were incubated at 37C for 16h in 0.5% (g/v) ammonium oxalate, and 1h at 85°C in 11% (v/v) peracetic acid. The resulting delignified AIR (dAIR) was washed three times with water and twice with acetone. dAIR powder was resuspended in water to a concentration of 10 mg ml^-1^.

### Oligosaccharide mass spectrometry profiling

Reaction mixtures (20 µl) contained 10 µl of 10 mg ml^-1^ resuspended dAIR, 1 µl of *Si*GH11 and 9 µl of water and were incubated for 16h at 30 °C. For subsequent treatment, the sample was centrifuged, the supernatant harvested and incubated with 1 µl water, EV, *Si*AXE or 250 mM NaOH and incubated for 16h at 30 °C, prior to usage of 1 µl for MALDI-ToF as previously described with small modifications ^59^. A layer of matrix crystals was formed onto a MALDI target plate by vacuum-drying 2 µl of a solution of 10 mg ml^-1^ 2,5-dihydroxybenzoic acid in 50% (v/v) acetonitrile and 10 mM NaCl. Reaction supernatants (1 µl) were deposited on top of the matrix crystals and dried under vacuum. Positive ion MALDI mass spectra were acquired with a RapifleX MALDI-ToF (Bruker) instrument operating in reflectron mode with an acceleration voltage of 20 kV. In each spectrum, 10000 shots were averaged. Relative abundance of xylan oligosaccharides with different acetyl substituents was calculated based on the area of the corresponding ions in the MALDI-ToF spectra.

### Size-exclusion chromatography of *Si*GH11-digested barley dAIR

Size exclusion chromatography (SEC) was carried out to separate *Si*GH11-released oligosaccharides according to their size. Barley dAIR slurry (10 mg ml^-1^) was incubated with *Si*GH11 for 16h, centrifuged and the supernatant split into two. One half was treated with 250 mM NaOH to obtain deacetylated barley XOS and the other half left untreated. SEC analysis was carried out on a NGC Scout 10 chromatography system (Bio-Rad), equipped with a Superdex Peptide Column 10/300 GL (Cytiva). The eluent (degassed water) was pumped at a flow rate of 0.5 ml min^-1^. The analytical columns were calibrated with dextran standards (2.5 and 6kDa; Roth). In total, fifteen fractions were collected with 1.5 ml volume each and dried in a Concentrator plus SpeedVac (Eppendorf). The fractions were resuspended in 100 µl water and an aliquot (1 µl) used for MALDI-ToF analysis. 25 µl of the remaining fractions were tested for their ability to induce apoplastic ROS production in barley roots.

### Determination of monosaccharide composition and cell wall bound acetate in barley root cell wall

To test whether *Si*AXE acts directly on the complex CW polymer or on soluble oligosaccharides, barley dAIR slurry (10 mg ml^-1^) was incubated with *Si*GH11 alone or in combination with *Si*AXE for 16h. Then, soluble oligosaccharides were harvested by centrifugation and hydrolyzed hydrolyzed in 2 M trifluoroacetic acid as previously ^57^. Neutral monosaccharides and uronic acids content were determined using high-performance anion-exchange chromatography (HPAEC) in an Azura HPAEC system (Knauer) equipped with a Dionex CarboPac Pa20 column (Thermo Fisher Scientific) and a pulsed amperometric detector (PAD). The remaining dAIR pellet was washed and determination of total wall acetate content was performed as previously described ^60^. Briefly, wall material was incubated in 0.25 M NaOH for 1h. Supernatants were neutralized with HCl and the acetic acid content determined with the Acetic Acid Assay Kit (Megazyme; #K-ACET).

### ROS accumulation assay

ROS assays were performed as described previously with some modifications ^52^. Roots of germinated barley seedlings (4 d) were separated and cut into pieces of 0.5 cm length. Four root pieces were transferred to each well of a 96-well microtiter plate (white, flat bottom) containing 200 μl of MQ-H_2_O. The plate was incubated ON at 21°C for recovery in the dark. The next day, the water was replaced with water containing 20 µM LO-12 and 20 µg ml^-1^ HRP. After 15 min incubation in the dark, 100 µl two-fold concentrated elicitor solution was added to each well and chemiluminescence was measured immediately using a TECAN SPARK 10M microplate reader over all wells for 2h with an integration time of 450 msec. As elicitors, xylose, xylobiose, xylotriose, xylotetraose, xylopentaose, xylohexaose were used at a final concentrations of 450 μM, alone or in an equimolar mixture. In addition, samples from enzymatic digests were used as elicitors: As test of xylanase activity, *Si*GH11-digested Xyl1-Xyl6were used at final concentration corresponding to 450 μM substrate. Furthermore, naturally occurring CW-released and deacetylated oligosaccharides from barley dAIR were tested. For this, 300 μl barley dAIR slurry (10 mg ml^-1^) was mixed with 150 μl *Si*GH11 and incubated for 16h at 28°C under continuous shaking, centrifuged and the supernatant split into separate reactions of 100 μl and further incubated with 100 μl *Si*AXE, *Si*AXE^S226G^, water or 25 mM NaOH for 16h. The reactions were terminated at 95°C for 5 min, centrifuged again and the supernatant diluted 1:10 and used as elicitor (1:20 v/v final dilution).

### Bioinformatical analysis

All analyses were performed using R Statistical Software version 4.2.2.

### Weighted Co-Expression Network Analysis (WCNA)

RNA-seq data of *Serendipita indica* (*Si*) grown on 1/10 PNM medium (Mock) or colonizing the roots of *Hordeum vulgare* (*Hv*) or *Brachypodium distachyon* (*Bd*) was analysed with a WCNA approach in R ^61^. For this purpose, raw count values for the samples listed in supplementary table 4 deposited under the proposal number 505829 (Zuccaro 2020) were downloaded from the U.S. Department of Energy Joint Genome Institute (JGI) genome portal (https://genome.jgi.doe.gov/portal/). The raw counts were normalized using the a variance stabilizing vst() transformation from the DESeq2 package version 1.38.3 in R ^62^. Subsequently, samples with < 500.000 reads and genes with < 10 raw read counts in ώ10 % were excluded from further analysis. In consequence, expression values for 10,658 of 11,767 annotated *Si* genes (90 %) were used as input for the WCNA (version 1.72). The blockwiseModules() function was run with the following parameters: power = 12, networkType = “signed hybrid”, minModuleSize = 100, mergeCutHeight = 0.15. In total, 8,657 genes were assigned to seven different modules whereas 2,011 genes remained unassigned (M0). Compared to the whole *Si* genome as background, two modules (M4 and M7) were >2-fold enriched for genes encoding for putatively secreted proteins as identified with SignalP version 6 ^63^. The connections between the genes in these modules were visualized with Cytoscape version 3.10.2 ^64^. Furthermore, a Gene Ontology (GO) enrichment analysis was performed on these genes using the enricher() function of ClusterProfiler version 4.6.0 ^65^. Functional annotations of *Si* genes were downloaded from Mycocosm (https://mycocosm.jgi.doe.gov/mycocosm/home). The results of the monocot-specific WCNA were compared to a WCNA generated based on RNA-seq data of *Si* during the colonization of the dicot host *Arabidopsis thaliana* (*At*).

### Differential gene expression analysis

Significant differentially expressed genes (in comparison to mock samples were identified for all timepoints using the DESeq2 package version 1.38.3 in R ^62^. For the generation of heatmaps, log_2_FC values were visualized using the pheatmap package version 1.0.12.

## Data availability

All data supporting the findings of this study are available within the article and supplementary material.

## Acknowledgements

MP, RE, PS, LA, and AZ acknowledge support from the Cluster of Excellence on Plant Sciences (CEPLAS), funded by the Deutsche Forschungsgemeinschaft (DFG, German Research Foundation) under Germany’s Excellence Strategy – EXC 2048/1, Project ID 390686111. AZ and AE acknowledge support from the SPP2125 DECRyPT.

## Author contributions

MB and AZ conceived and designed the research. MB and AZ wrote the manuscript. MB, VRG, RE, PS, and ABE performed the experiments. MB, VRG, RE, PS, MP, and ABE analyzed the data. LA conducted the bioinformatic analysis of co-regulated genes. AZ and MP provided funding for the experiments. All authors contributed to manuscript editing and approved the final version.

---

**Extended Data Fig. 1.**
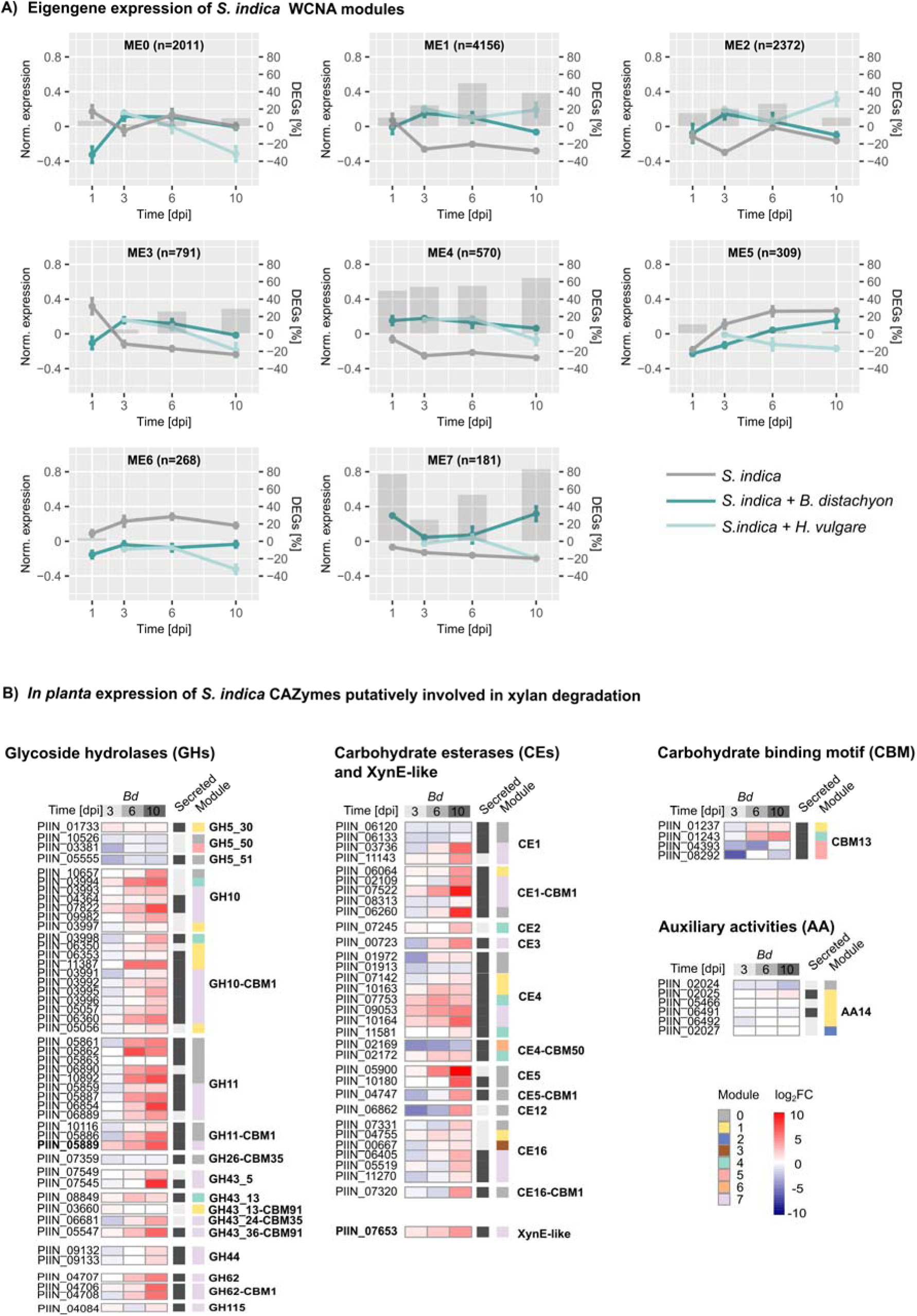
Eigengene expression of *S. indica* WCNA modules and expression of xylan-targeting CAZymes during *B. distachyon* colonization. **A)** Eigengene expression of *S*. *indica* WCNA modules 0-7. Shown is the expression of representative genes (Eigengenes) for co-expressed modules. Module 7 comprises 181 genes that are highest expressed at late stage of monocot *B. distachyon* (*Bd*) host colonization. Gene expression data are from *S. indica* from axenic culture (grey) or colonizing the monocot hosts *Bd* (dark green) or *H. vulgare* (light green). B) Gene expression profiles of *S. indica* carbohydrate-active enzymes (CAZymes) including glycoside hydrolases (GH), carbohydrate esterases (CE), CBM13-containing proteins, and auxiliary activity (AA) enzymes. Heatmaps display mean expression (log2FC) of genes implicated in xylan hydrolysis, alongside their WCNA module assignments. RNA-Seq data were obtained from *S. indica* colonizing the monocot host (*Bd*), at 3, 6, and 10 days post-inoculation (dpi) from previously published data 21. Note: CAZyme annotations are based on CAZyDB (http://www.cazy.org/), dbCAN3-HMMER, dbCAN3-sub ^4–5^, and manual curation by subfamily classification (see Table S1). *SiGH11* (PIIN_05889) and the XynE-like esterase *SiAXE* (PIIN_07653) are highlighted in bold.

**Extended Data Fig. 2.**
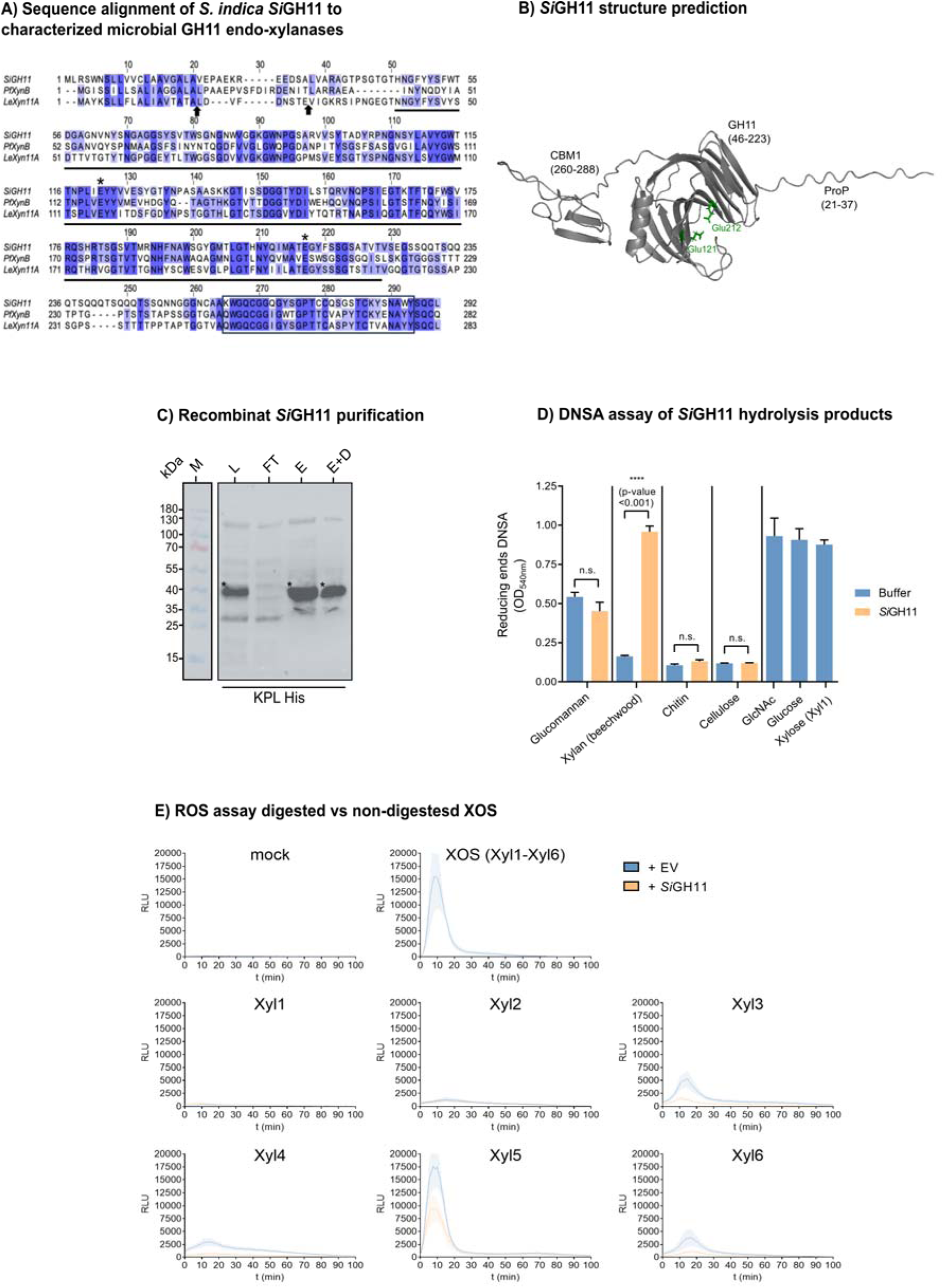
*Si*GH11 has structural similarity with functionally characterized microbial GH11 endo-xylanases. **A)** Protein of *Si*GH11 with characterized GH11 xylanases from *Penicillium funiculosum* (Q8J0K5.1) and *Lentinula eodes* (Q96UV7). Identical and conserved amino acids across sequences are color coded in dark and light blue, respectively. The position of the predicted cleavage sites for the signal peptide (SP) and propeptide (ProP) are indicated by arrows. The catalytic glutamic acid residues E121 (nucleophile) and E212 (acid/base catalyst) within the GH11 domain (underlined) are indicated by asterisks. The CBM1 domain is indicated by a black box. **B)** AlphaFold prediction of *Si*GH11. The N–terminal signal peptide (1-20) was removed prior to structure prediction. *Si*GH11 contains a propeptide (ProP) at amino acid position 21-37, a glycoside hydrolase 11 (GH11) domain at position 46-223 and a carbohydrate binding motif 1 (CBM1) at position 260-288. Core catalytic residues (Glu121 and Glu212) are colored green. **C)** Recombinant protein production and extraction of *S. indica* GH11. Protein bands corresponding to *Si*GH11 are indicated by asterisk. His-tagged protein was detected by immunoblotting with KPL His detector. Abbreviations: L=lysis, FT=flow-through, E=Elution, E+D=Elution+dialysis. **D)** *Si*GH11 activity was tested on glucomannan, xylan (beechwod), chitin, cellulose and acetyl-xylan (birchwood). As control, the cell wall monomers N-acetyl Glucosamin (GlcNAc), glucose and xylose (Xyl1) were used. After incubation for 16h, the enzyme reactions were centrifuged and the supernatant containing soluble oligosaccharides was used for DNSA assay to quantify reducing end sugars. Hydrolytic activity of *Si*GH11 on beechwood xylan resulted in increased abundance of reducing ends; i.e the xylooligosaccharide hydrolytic products. Asterisks indicate significant differences based on Student’s t-test: ns, not significant; ***, p ≤ 0.001. **E)** Reactive oxygen species (ROS) burst in barley root segments triggered by XOS (Xyl1–Xyl6) before and after *Si*GH11 treatment (16 h incubation). *Si*GH11 digestion reduces ROS induction by longer XOS (Xyl3–Xyl6). A mixture of Xyl1–Xyl6 served as a positive control. The graphs show mean ± s.e.m of four wells each containing four root segments. The same data are plotted as area under the curve (AUC) in figure 3C.

**Extended Data Fig. 3.**
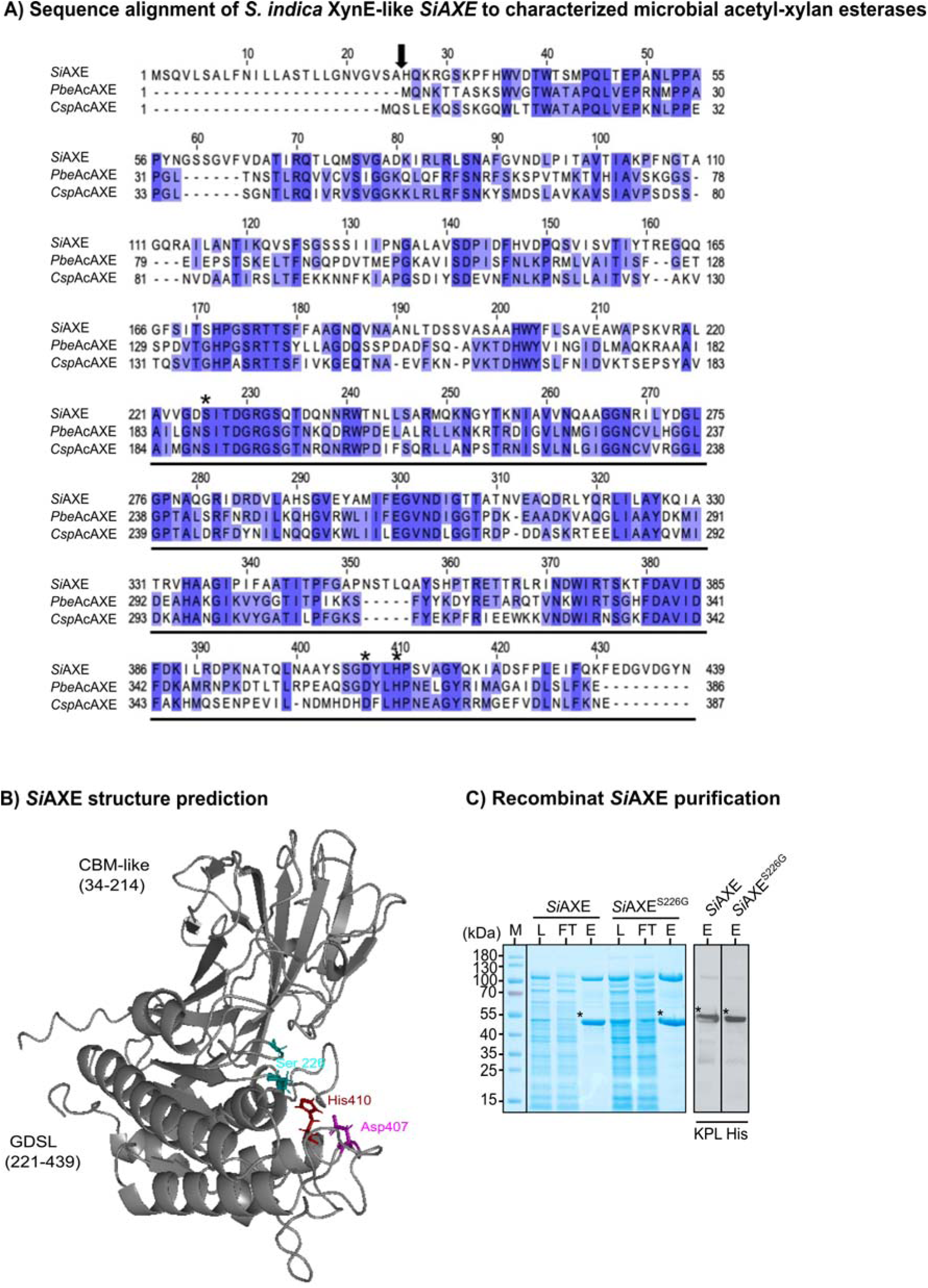
*Si*AXE has structural similarity with functionally characterized microbial acetyl-xylan esterases. **A)** Multiple sequence alignment of *Si*AXE with characterized acetyl xylan esterases from *Prolixibacter bellariivorans* (A0A5M4AV20) and *Chryseobacterium sp. YR480* (A0A1M6XU10). Identical and conserved amino acids across sequences are color coded in dark and light blue, respectively. The position of the predicted cleavage site for the *Si*AXE signal peptide (SP) is indicated by a black arrow. The predicted catalytic Ser_226_, Asp_407_ and His_410_ residues within the GDSL domain (underlined) are indicated by black asterisks. **B)** AlphaFold structure prediction of *Si*AXE. *Si*AXE has a predicted CBM-like domain at amino acid position 34-214 and a Lipase_GDSL_2 domain at position 221-439. *Si*AXE belongs to the SGNH hydrolase superfamily with the core residues Ser226 (teal), Asp407 (pink) and His410 (red). **C)** Recombinant protein production and extraction of *Si*AXE and the catalytic mutant SiAXE^S226G^. Protein bands corresponding to *Si*AXE are indicated by asterisks. His-tagged protein was detected by immunoblotting with KPL His detector. Abbreviations: L=lysis, FT=flow-through, E=Elution, E+D=Elution+dialysis.

**Extended Data Fig. 4.**
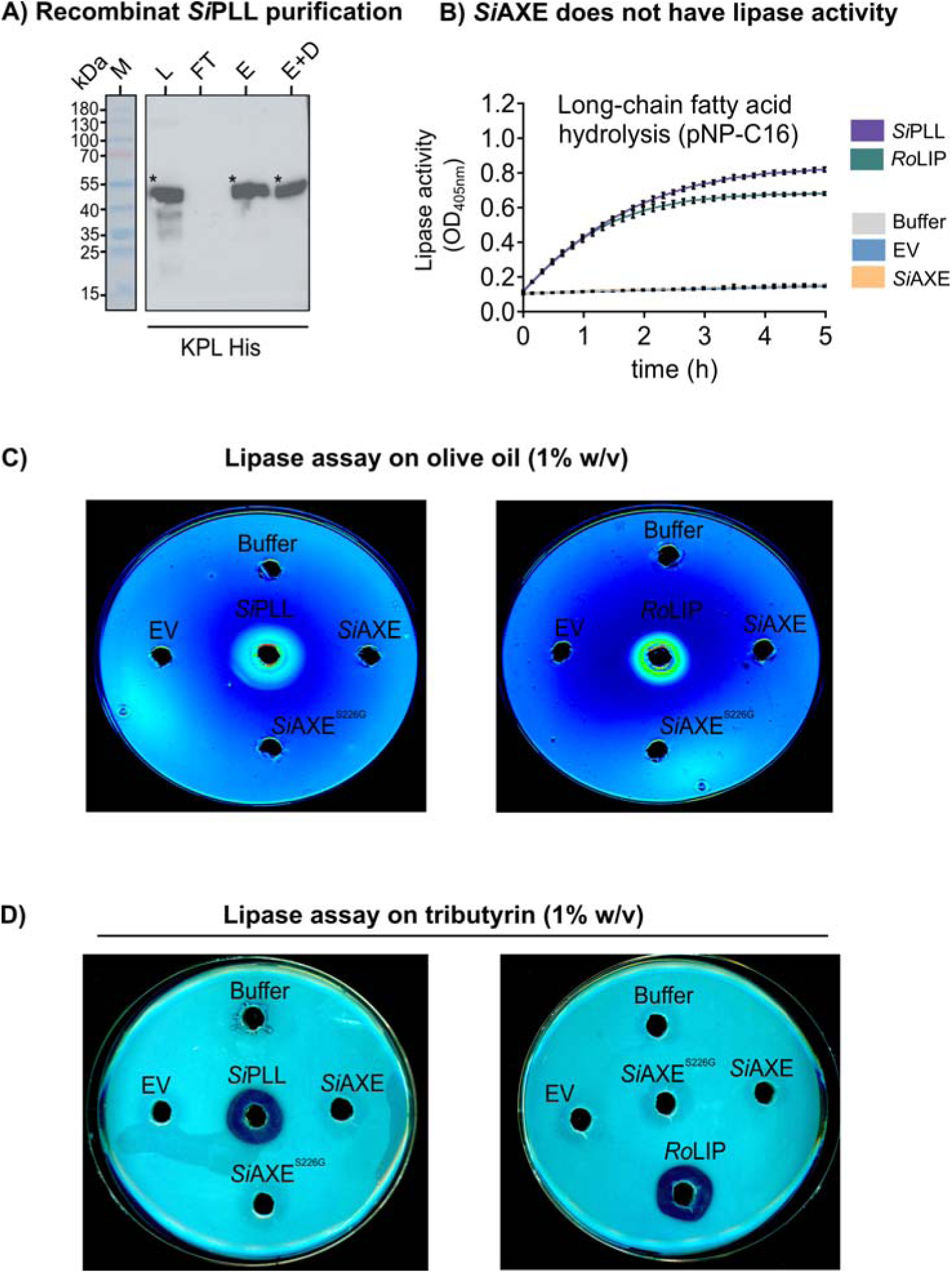
*Si*AXE has no lipase activity. **A)** Recombinant protein production and extraction of *Si*PLL used as positive control for the lipase activity assays. Protein bands corresponding to *Si*PLL are indicated by asterisks. His-tagged protein was detected by immunoblotting with KPL His detector. Abbreviations: L=lysis, FT=flow-through, E=Elution, E+D=Elution+dialysis. **B)** Lipase activity of recombinant protein with the long-chain artifical substrate *para*-nitrophenyl-palmitate (pNP-C16). Yellow color indicates shift in absorbance at OD_405nm_ representative of long-chain acyl ester hydrolysis. In contrast to *Si*AXE, the promiscous lipase *Si*PLL from *S. indica* and the commercial lipase from *Rhizopus oryzae* (*Ro*LIP; Merck #62305) are highly active with the lipid substrate pNP-C16. **C)** Rhodamine B plate assay of lipase activity on olive oil-containing plates. Lipid hydrolysis of the olive oil lipids is seen by the formation of a halo around the enzyme solution after excitation at 545 nm. **D)** Plate assay of lipase activity on tributyrin-containing plates. Hydrolysis of the tributyrin esters is seen by the formation of a halo where the enzyme solution was applied. Lipase activity on tributyrin is visible for *Si*PLL and *Ro*LIP.

**Extended Data Fig. 5.**
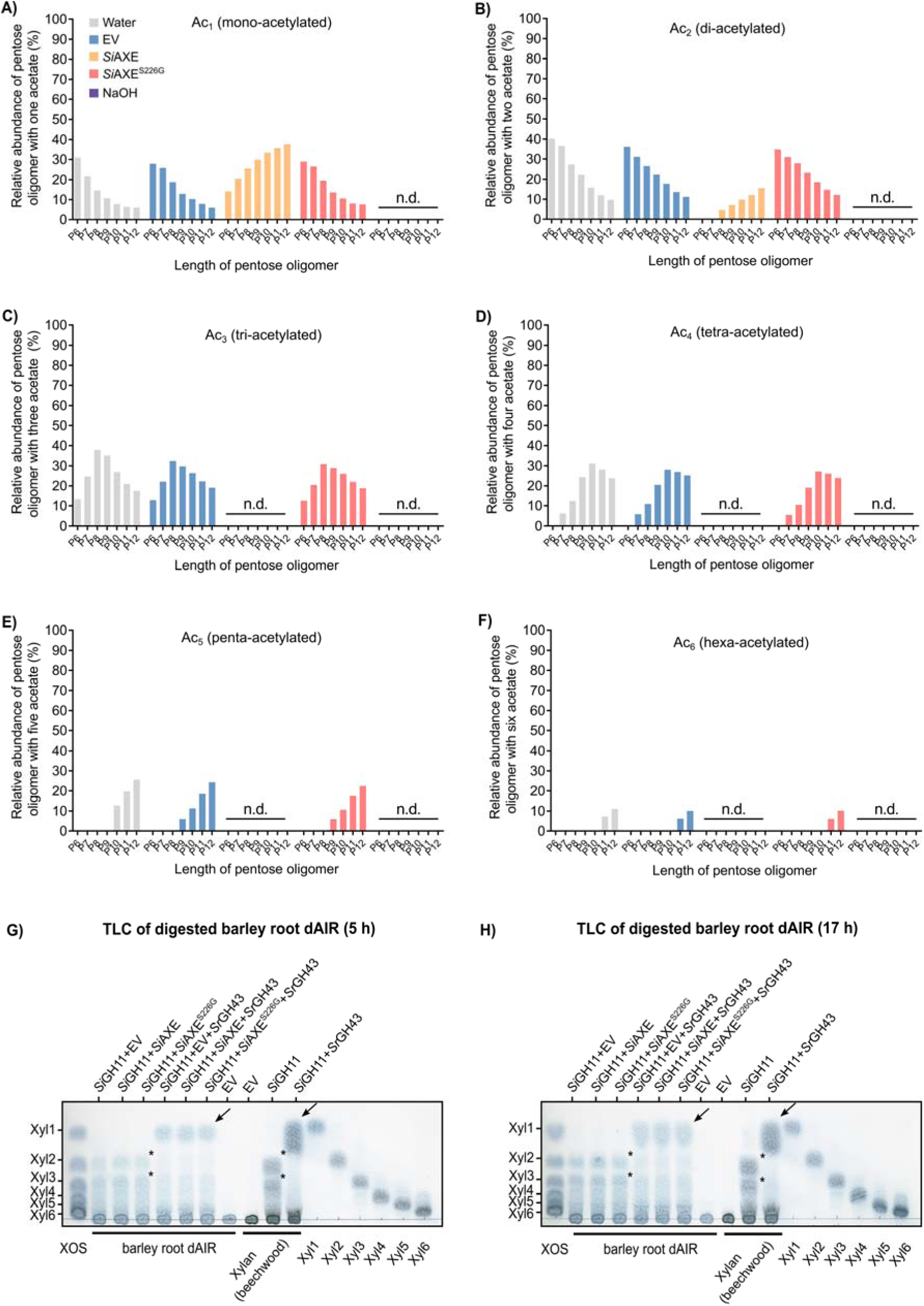
*Si*GH11-released acetylated xylooligosaccharides from barley root hemicellulose are deacetylated by *S*iAXE and hydrolyzed by a GH43 exo-xylanase. **A)** Barley root dAIR was incubated with *Si*GH11 and the released soluble oligomers were subsequently treated with EV, *Si*AXE, *Si*AXE^S226G^ or NaOH and analyzed by MALDI-ToF (see Figure 4D-F). The length of the pentose oligomers (P6-P12) is plotted on the x-axis and the relative abundance of the acetylation of the respective pentose oligomer is plotted on the y-axis. Incubation with *Si*AXE or NaOH decreased abundance of single-acetylated pentose hexamers (P_6_Ac_1_ and P_7_Ac_1_). NaOH treatment was stronger and led to absence of P6-P12Ac_1_, while abundance of P8-P12-Ac_1_ was increased after *Si*AXE treatment. **B)** Incubation with *Si*AXE or NaOH decreased abundance of di-acetylated pentose oligomers. NaOH treatment was stronger and led to absence of P6-P12Ac_2_, while *Si*AXE treatment led to absence of P6Ac_2_ and decrease of P7-P12Ac_2_. **C)** Incubation with *Si*AXE or NaOH decreased abundance of tri-acetylated pentose oligomers (P6-P12Ac_3_). **D)** Incubation with *Si*AXE or NaOH decreased abundance of tetra-acetylated pentose oligomers (P6-P12Ac_4_). **E)** Incubation with *Si*AXE or NaOH decreased abundance of penta-acetylated pentose oligomers (P6-P12Ac_5_). **F)** Incubation with *Si*AXE or NaOH decreased abundance of hexa-acetylated pentose oligomers P6-P12Ac_6_). **G-H)** TLC of soluble hydrolysis products containing oligosaccharides obtained from barley dAIR digest by *Si*GH11, *Si*AXE and *Sr*GH43 alone or in combination after 5h (G) and 17 h (H) (see Figure 4G). The enzyme combinations used are indicated at the top of the plate. The first lane is a mixture of all six XOS that are also individually spotted on the last six lanes, as indicated below the plate. Xylobiose and xylotriose released from barley dAIR by SiGH11 are indicated by an asterisk. Xylose produced by *Sr*GH43 exo-xylanase is indicated by an arrow. As additional control for xylanase activity, xylan (beechwood) was used as substrate.

**Extended Data Fig. 6.**
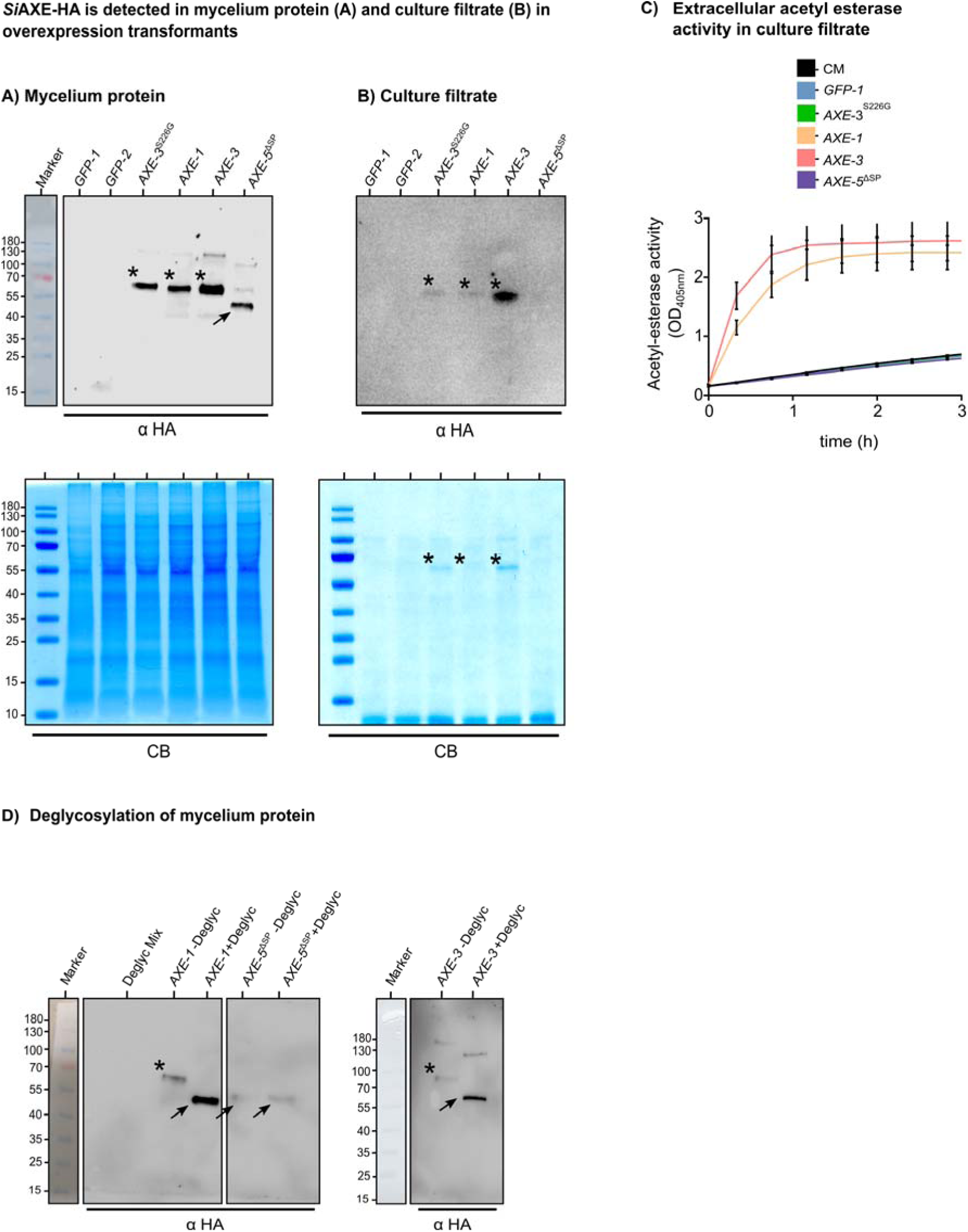
*Si*AXE is secreted and confers extracellular acetyl-esterase activity. **A)** Western blot using α-HA antibody with protein extracted from mycelium grown in axenic culture. The immunoblot signal between 55-70 kDA, indicated by the asterisk, corresponds to glycosylated *Si*AXE-HA in transformants expressing full-lenght *SiAXE* (*AXE-1* and *AXE-3*), and the catalytic inactive version (*AXE-3*^S226G^). In the transformants with deleted N-terminal signal peptide (*AXE-5*^ΔSP^) an immunoblot signal at ∼55 kDA, corresponding to deglycosylated *Si*AXE-HA is indicated by the arrow. **B)** Western blot using α-HA antibody with culture filtrate from *S. indica* grown in axenic culture. The band marked by the asterisk corresponds to glycosylated *SiAXE*-HA. In the transformants with deleted N-terminal signal peptide (*AXE-5*^ΔSP^) no signal is detected, consistent with absence of secretion into the culture medium. **C)** Extracellular acetyl-esterase activity with artificial substrate *para*-nitrophenyl acetate (pNP-C2) in *S. indica* transformants. Yellow color is indicative of acetyl-ester hydrolysis and results in a shift of absorbance at OD_405nm_. In culture filtrate from transformants expressing full-lenght *SiAXE* (*AXE-1* and *AXE-3*), high extracellular acetyl-esterase activity is detected, while it is absent in CM medium and in culture filtrate from *GFP-1*, *AXE-3*^S226G^ and *AXE-5*^ΔSP^. **D)** Deglycosylation of protein extracted from mycelium grown in axenic culture from transformants *AXE-1*, *AXE-5*^ΔSP^ and *AXE-3*. The protein extracts were either treated with protein deglycosylation mix II (NEB; #P6044S) (+Deglyc) or with buffer only (-Deglyc) for 16 h at 22 °C prior to application to SDS-PAGE and immunoblotting using α-HA antibody. Incubation with the deglycosylation mix leads to shift of the *SiAXE*-HA band in *AXE-1* and *AXE-3* from ∼70 kDa (marked by asterisk) to a band with a size of∼50-55 kDA (marked by arrow). The enzymatically deglycosylated *Si*AXE-HA band in *AXE-1* and *AXE-3* (+Deglyc) is comparable to that obtained in *AXE-5*^ΔSP^, which indicates that the removal of the signal peptide in *AXE-5*^ΔSP^ leads to the absence of deglycosylation.

**Extended Data Fig. 7.**
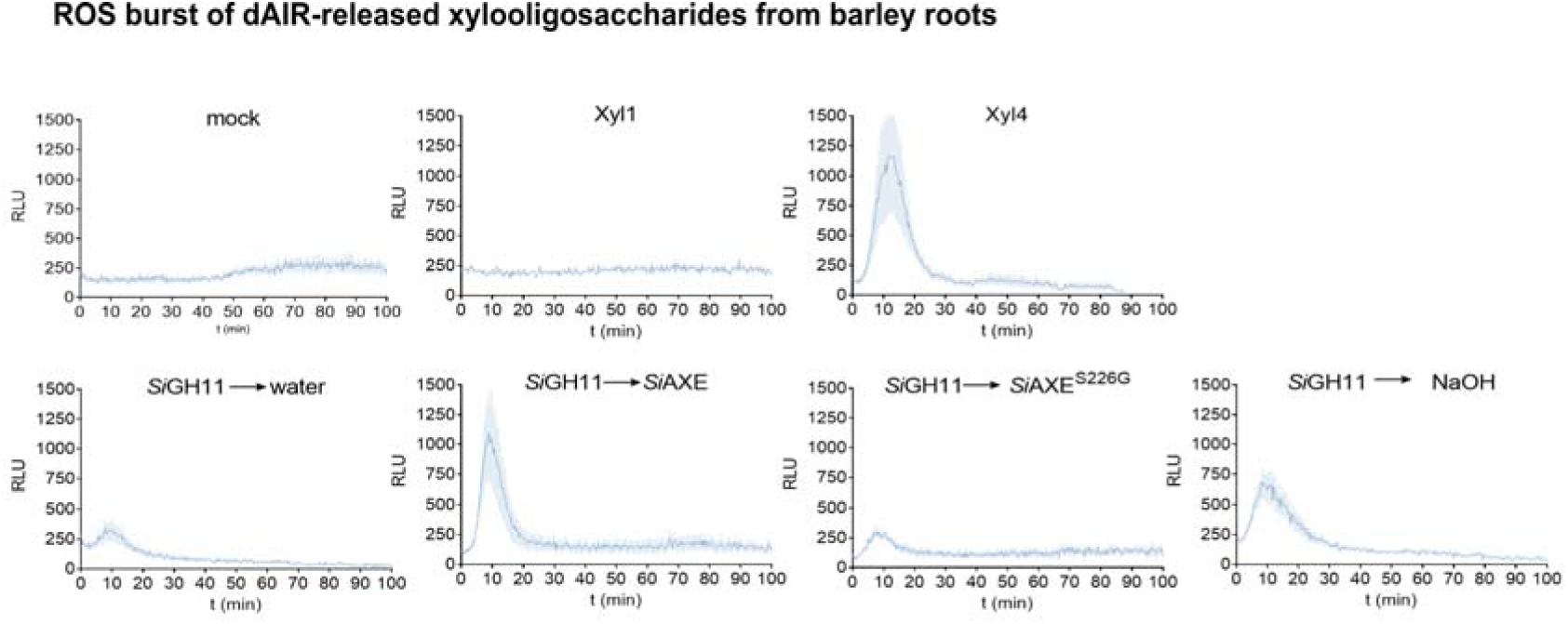
Accumulation of apoplastic ROS in barley roots upon elicitor treatment with xylooligosaccharides released from barley dAIR. ROS burst assay in barley root fragments treated with soluble xylooligosaccharides generated by enzymatic digestion of barley dAIR. Supernatants of *Si*GH11 digests were harvested and further treated with *Si*AXE, controls or NaOH prior to application as depicted schematically in Figure 4D. The graphs show mean ± s.e.m of four wells each containing four root segments. The same data are plotted as area under the curve (AUC) in figure 5E.

**Extended Data Fig. 8.**
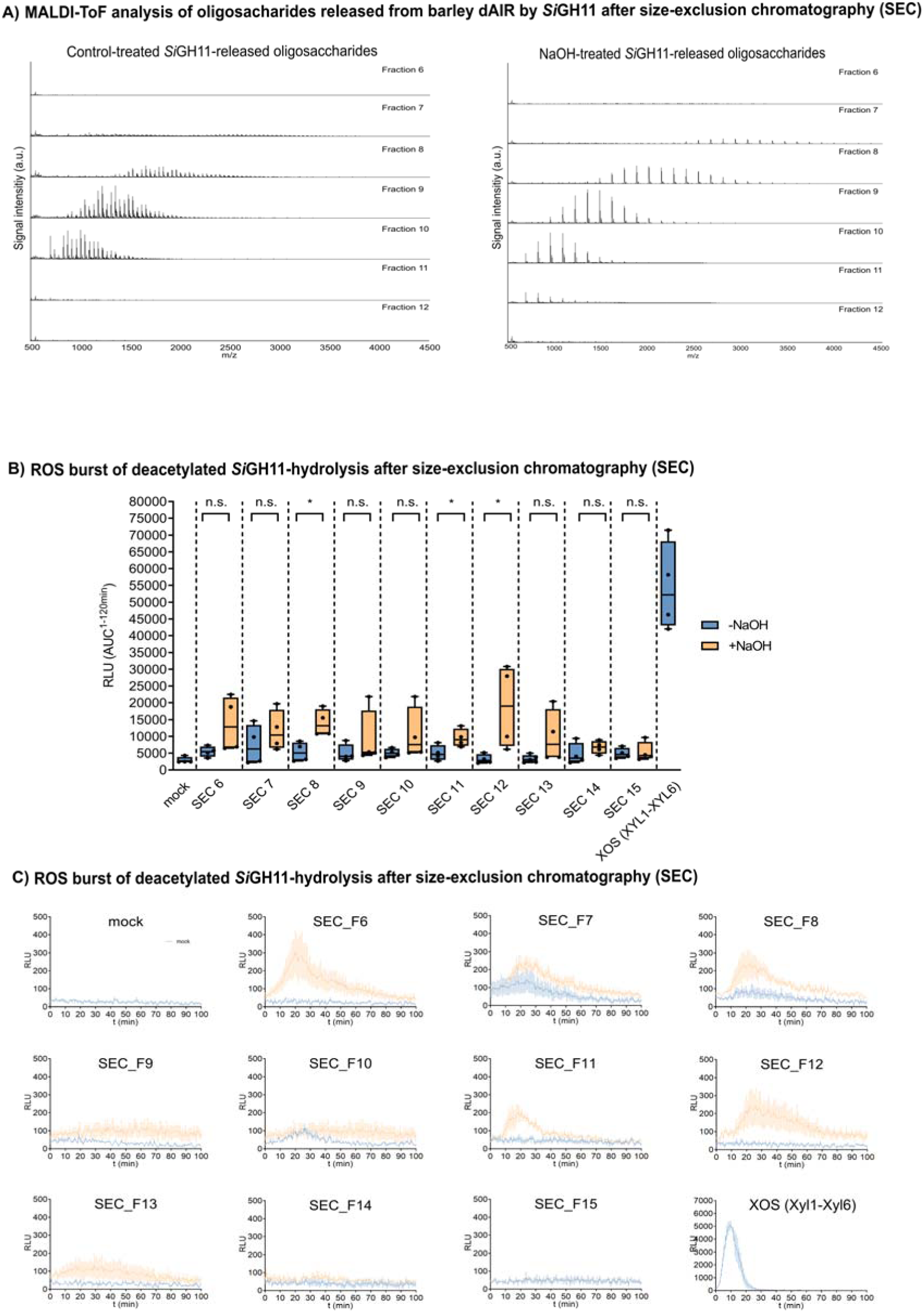
Deacetylated xylooligosaccharides of various lenght derived from barley dAIR elicit ROS responses. **A)** MALDI-ToF spectra of oligosaccharides released from barley dAIR after *Si*GH11 hydrolysis. The soluble oligosaccharides in the supernatant from *Si*GH11 hydrolysis of dAIR were treated with NaOH for chemical deacetylation or water as control. Then, the total hydrolysate was subjected to size-exclusion chromatography (SEC) to separate xylooligosaccharides (XOS) of various length. In total, 15 fractions were collected and analyzed by MALDI-ToF to confirm the release of XOS. The degree of polymerization (DP) eluted from the column in the individual fractions are as follows: Fractions 6-8 (>DP20), fraction 9 (DP11-20), fraction 10 (DP7-15), fractions 11-13 (DP<7). **B)** ROS burst assay in barley root fragments treated with the respective SEC fractions shown in A). Apoplastic ROS was measured by luminol-HRP chemiluminescence and quantified as area under the curve (AUC). See C for kinetics. As positive control for XOS-induced ROS accumulation, a mixture of XOS (Xyl1-Xyl6) was used. As negative control, the roots were treated only with water (mock). Asterisks indicate significant differences based on Student’s t-test: ns, not significant; *, p ≤ 0.05. RLU= Relative light units. **C)** The graphs show mean ± s.e.m. of four wells each containing four root segments as plotted as AUC in B).

**Extended Data Fig. 9.**
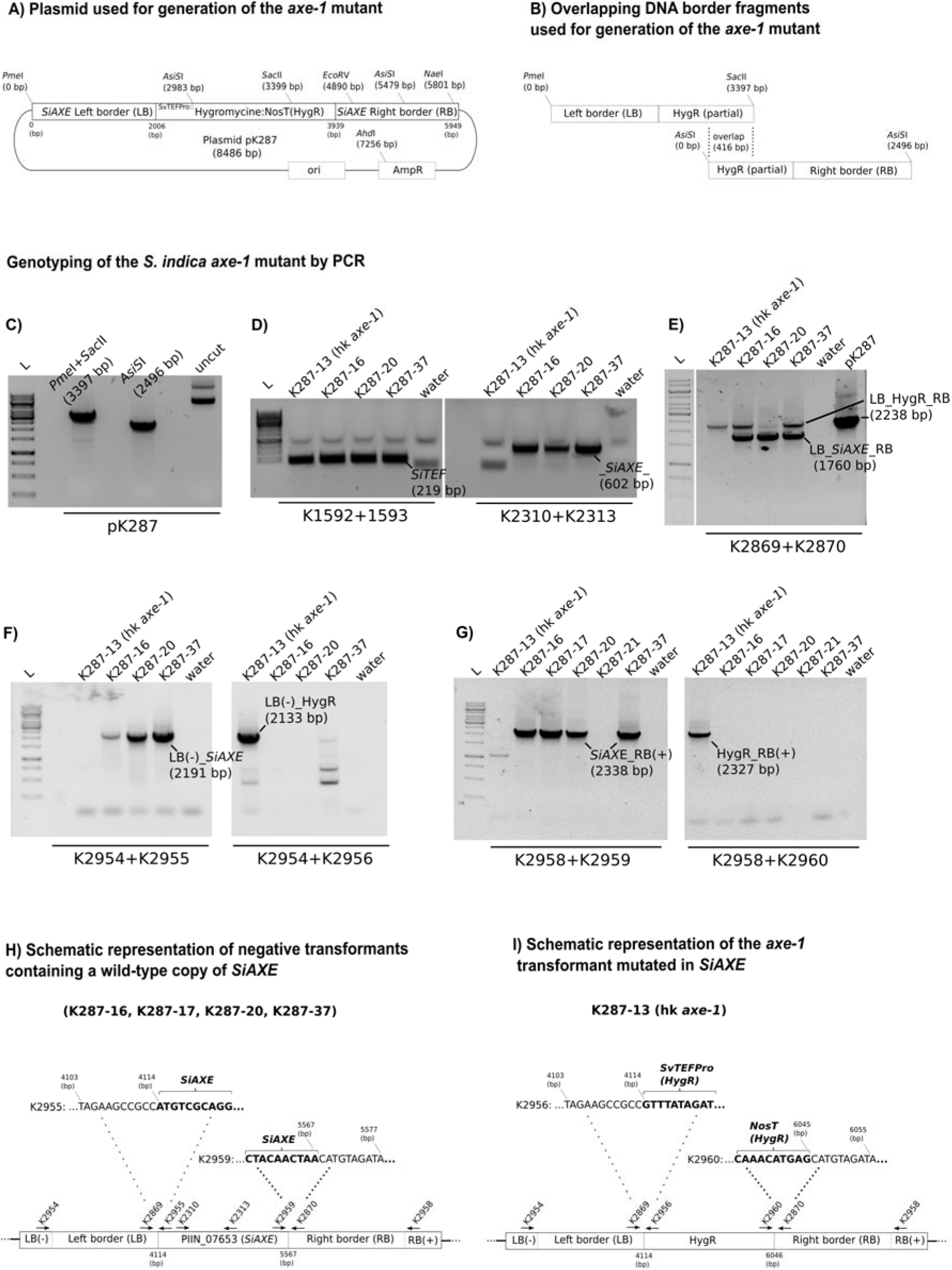
Generation of *axe-1* mutant in *S. indica* homokaryotic strain (hk#58 WT) **A)** Schematic representation of the plasmid (pK287) used for homologous recombination to replace the *SiAXE* (PIIN_07653) gene locus (Ensembl Fungi genome assembly ASM31354v1, position 4114-5567 bp on SuperContig PIRI_contig_0252; forward strand) in the *S. indica* genome. For generation of pK287, 2006 bp upstream (left border) and 2010 bp downstream (right border) of the *SiAXE* gene were amplified from *S. indica* genomic DNA via PCR and cloned in between a hygromycine resistance cassete (HygR) consisting of the *TEF* promotor from *Serendipita vermifera* (SvTEFPro), the hygromycine gene and the Nos terminator (NosT). **B)** Schematic representation of the linear DNA fragments used for protoplast-mediated transformation. To generate overlapping fragments for homologous recominbation, the pK287 plasmid was digested with *Pme*I and *Sac*II to generate the left border (LB) and with *Asi*SI to generate the right border (RB). These fragments were excised from agarose gel and used for transformation of homokaryotic *S. indica* strain hk#58 WT. **C)** Confirmation of the linear LB and RB fragments, the LB (*Pme*I+*Sac*II digest) is 3397 bp, the RB (*AsiI*SI digest) is 2496 bp and the uncut plasmid pK287 is 8486 bp in lenght. **D-G)** Genotyping of the *axe-*1 mutant via PCR. Shown here is a selection of the transformants that are resistant to hygromycine but have wild-type copies of *SiAXE* (K287-16, K287-17, K287-20, K287-21, K287-37) and therefore serve as genetic background controls for the *axe-1* mutant (K287-13). **D)** The fungal housekeeping gene *SiTEF* was successfully amplified with primers K1592+K1593 in all samples, while amplification of *SiAXE* with primers K2310+K2313 did not result in a PCR product in the *axe-1* mutant. **E-G)** The binding sites of the primers used to verify the replacement are indicated schematically in (H) and (I) for the wild-type copy of SiAXE and the *axe-1* mutant, respectively. The primer pairs used for amplification of the various PCR products in D-G are indicated below each gel image. The name and length of the amplified PCR products in are indicated in each gel image. PCR with primers K2869+K2870 in E) resulted in the two PCR products LB_HygR_RB (2238 bp) and LB_AXE_RB (1760 bp) in the genetic background controls K287-16, K287-20 and K287-37. In *axe-1* (K287-13), only the first product (LB_HygR_RB; 2238 bp) was amplified, corresponding to the PCR product obtained with the vector pK287. This indicates that there is integration of the HygR cassette in the genetic background control, while the *SiAXE* genomic locus remains intact. **H-I)** Sequencing of the indicated PCR products amplifying the junction between the *SiAXE* genomic locus (in H), the HygR locus (in I) and the LB-RB border sequences, respectively, returned the indicated nucelotide sequence, confirming seamless replacement of the *SiAXE* locus by HygR in *axe-1* (I). The *SiAXE* sequence (in H) and the HygR sequence (in I), located in between LB-RB, are indicated in bold, respectively. Abbreviations: L= ladder, LB= left border, RB= right border, HygR= hygromycine resistance, LB(-)= region upstream of the 2000 bp LB, RB(+)= region downstream of the 2000 bp RB.

**Extended Data Fig. 10.**
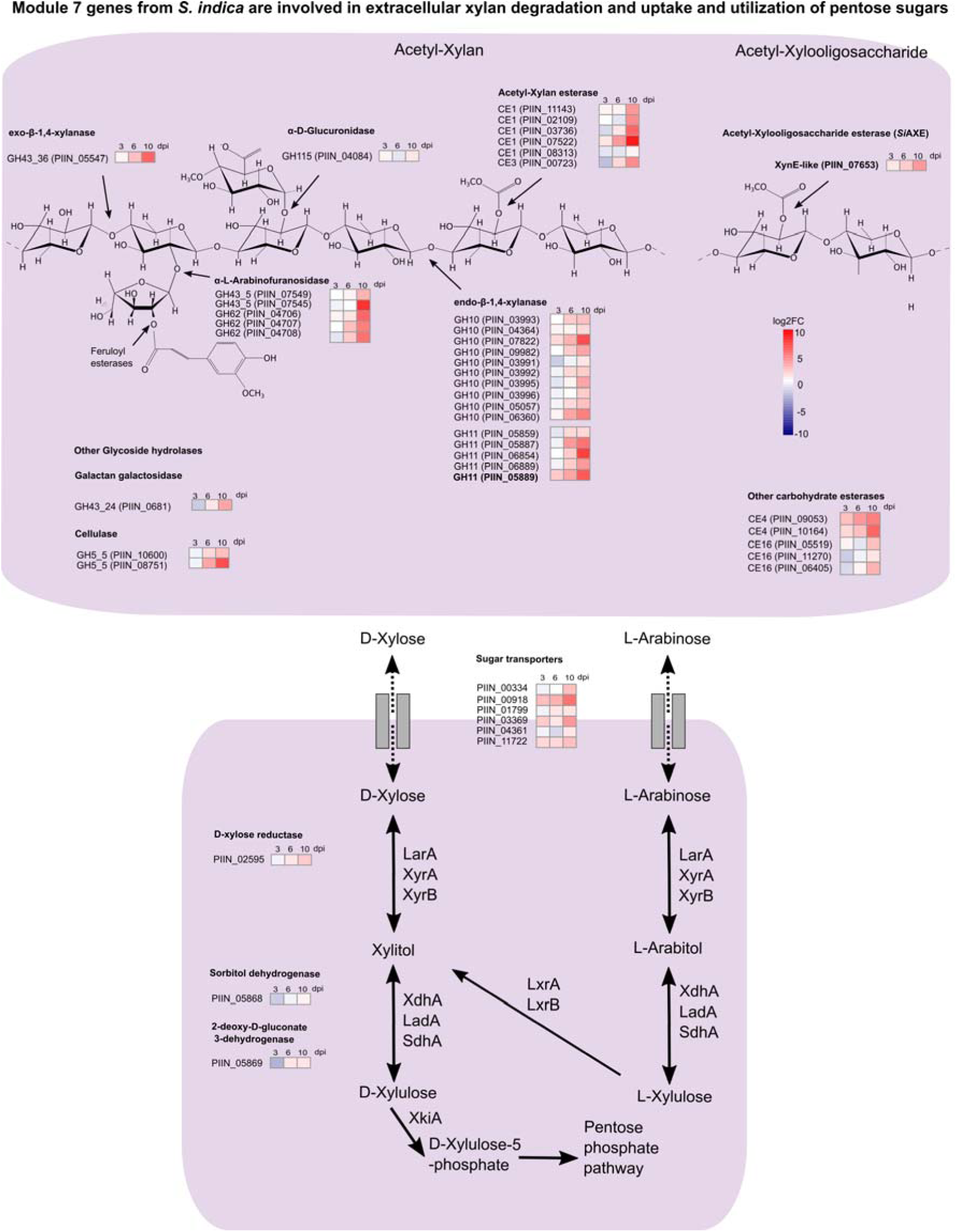
Extracellular xylan hydrolysis and uptake of pentose sugars by *S. indica* module 7 protein. **A)** Genes encoding extracellular xylanolytic enzymes from module 7 comprise putative α-L-Arabinofuranosidases from families GH43_5 and GH62, an α-D-Glucuronidase from family GH115, endo-β-1,4-xylanases from families GH10 and GH11, exo-β-1,4-xylanass from family GH43_36, acetyl-xylan esterases from families CE1 and CE3 and the XynE-like acetyl-xylooligosaccharide esterase *Si*AXE. Besides, module 7 comprises six putative sugar transporters that could function as pentose (xylose and arabinose) transporters and D-xylose reductase (PIIN_02595), sorbitol dehydrogenase (PIIN_05868), 2-deoxy-D-gluconate 3-dehydrogenase (PIIN_05869) that are involved in the pentose catabolic pathway in other fungi. The heatmaps show mean gene expression (log2FC) from *B. distachyon* colonized by *S. indica* at 3, 6 and 10 dpi (see Figure S1B). The annotation of pentose catabolic pathway enzymes is from *Aspergillus niger* ^51^.

