## Supplementary material for "Host-adapted enzymatic deconstruction of acetylated xylan enables mutualistic colonization of monocot roots": Table S4

**Supplementary Table 4**: Samples downloaded from the JGI genome portal for analysis. Due to low read numbers (< 500.000), the samples Zuccaro_RNA.P_HvSi1_1dpi, Zuccaro_RNA.P_HvSi2_1dpi and Zuccaro_RNA.P_HvSi3_1dpi were excluded from further analysis.

| **Sample Name** | **Number of reads** |
| --- | --- |
| Zuccaro_RNA.P_BdSi1_1dpi | 732157 |
| Zuccaro_RNA.P_BdSi2_1dpi | 697107 |
| Zuccaro_RNA.P_BdSi3_1dpi | 856796 |
| Zuccaro_RNA.P_BdSi1_3dpi | 6248471 |
| Zuccaro_RNA.P_BdSi2_3dpi | 6618380 |
| Zuccaro_RNA.P_BdSi3_3dpi | 3959427 |
| Zuccaro_RNA.P_BdSi1_6dpi | 4293067 |
| Zuccaro_RNA.P_BdSi2_6dpi | 3651198 |
| Zuccaro_RNA.P_BdSi3_6dpi | 4329712 |
| Zuccaro_RNA.P_BdSi1_10dpi | 4678533 |
| Zuccaro_RNA.P_BdSi2_10dpi | 5640310 |
| Zuccaro_RNA.P_BdSi3_10dpi | 6078590 |
| Zuccaro_RNA.P_HvSi1_1dpi | 476 |
| Zuccaro_RNA.P_HvSi2_1dpi | 691 |
| Zuccaro_RNA.P_HvSi3_1dpi | 288 |
| Zuccaro_RNA.P_HvSi1_3dpi | 2877428 |
| Zuccaro_RNA.P_HvSi2_3dpi | 2815821 |
| Zuccaro_RNA.P_HvSi3_3dpi | 2723926 |
| Zuccaro_RNA.P_HvSi1_6dpi | 1890284 |
| Zuccaro_RNA.P_HvSi2_6dpi | 1644781 |
| Zuccaro_RNA.P_HvSi3_6dpi | 2303279 |
| Zuccaro_RNA.P_HvSi1_10dpi | 9367151 |
| Zuccaro_RNA.P_HvSi2_10dpi | 5531302 |
| Zuccaro_RNA.P_HvSi3_10dpi | 8880937 |
| Zuccaro_RNA.P_Si1_1dpi | 5924010 |
| Zuccaro_RNA.P_Si2_1dpi | 11447260 |
| Zuccaro_RNA.P_Si3_1dpi | 14334607 |
| Zuccaro_RNA.P_Si1_3dpi | 22907988 |
| Zuccaro_RNA.P_Si2_3dpi | 12022301 |
| Zuccaro_RNA.P_Si3_3dpi | 17550090 |
| Zuccaro_RNA.P_Si1_6dpi | 20974919 |
| Zuccaro_RNA.P_Si2_6dpi | 10792651 |
| Zuccaro_RNA.P_Si3_6dpi | 6278384 |
| Zuccaro_RNA.P_Si1_10dpi | 9361226 |
| Zuccaro_RNA.P_Si2_10dpi | 7681926 |
| Zuccaro_RNA.P_Si3_10dpi | 9262132 |
